# Characterizing the viscoelastic properties of different fibroblasts in 2D and 3D collagen gels

**DOI:** 10.1101/2023.12.15.571880

**Authors:** Sandra Pérez-Domínguez, Héctor Sanz-Fraile, Laura Martínez Vidal, Massimo Alfano, Jorge Otero, Manfred Radmacher

**Affiliations:** Institute for Biophysics, University of Bremen, Bremen, Germany; Unit of Biophysics and Bioengineering, University of Barcelona, Cellular and respiratory biomechanics, Institute for Bioengineering of Catalonia, CIBER Respiratory diseases, Barcelona, Spain; Vita-Salute San Raffaele University, Milan, Italy; Division of Experimental Oncology/Unit of Urology, URI, IRCCS, Ospedale San Raffaele, Milan, Italy

**Keywords:** Dupuytren’s disease, collagen matrix, fibroblast, rheology, viscoelasticity

## Abstract

We assessed cell mechanical properties in both 2D and 3D environments employing compliant type I collagen matrices. Firstly, collagen gels of varying stiffness were prepared using a photocrosslinker to increase gel stiffness. Using methacrylic anhydride and UV light, a 10-fold increase in apparent Young’s modulus with respect to the soft collagen gel was achieved (0.2 kPa to 2 kPa). In addition, cells were plated onto the different collagen gels and hard Petri dishes (as a super stiff substrate) and their mechanical properties were evaluated. An increase in apparent Young’s modulus was observed in Dupuytren fibroblasts behavior when increasing substrate stiffness, supporting its myofibroblast phenotype (3.8 kPa to 5.2 kPa from soft collagen gels to hard Petri dishes). Secondly, gel’s mechanics, in which fibroblasts were embedded, were evaluated over time to assess cells contraction properties. Gel’s apparent Young’s modulus increased over time regardless of fibroblasts type and cells presented dendritic protrusions. Rheological properties of both cells and gels were extracted using AFM sweep frequency scheme and power law structural damping model for data analysis. As a summary, we have found that fibroblasts contractile properties, related to myofibroblast differentiation and development are highly influenced on the mechanical properties of the surrounding environment, being stiffer environments those that favor the increase in fibroblast mechanical tension.

## Introduction

Tissue is the biological organization of structurally and functionally similar cells and their extracellular matrix (ECM)^1^. In disease, it is important to study both cellular and ECM alterations to reach a better understanding of disease’s origin and progression^2–4^. In biophysics, studies at single cell level are commonly employed to assess cell mechanical changes, among other changes like adhesion or biochemical variations, in disease. AFM has been used to compare mechanical alterations between healthy and cancerous or pathological cells^5^, opening a new window supporting the potential of studying cell mechanics with AFM as a diagnostic marker. However, it is well known that cells react and respond to their surrounding environment; therefore, it is important to specify in which conditions measurements were performed. Several progresses in the topic showed that cancerous cells being softer than healthy cells could not be taken as a general statement. Healthy cells were found to modify its stiffness to the stiffness of the support in contrast to cancerous cells, which appear to be less sensitive to changes in substrate mechanical properties^6^. Currently, artificial and natural hydrogels are becoming popular as 2D scaffolds to plate cells and assess cell behavior under specific conditions. Polyacrylamide (PA) gels are widely used due to their capacity of being modulated to achieve large range variability in stiffness^7–9^. Artificial hydrogels made of natural polymers such as collagen, gelatin, hyaluronic acid, alginate, and dextrin are also employed to measure cell mechanics on and in compliant substrates. These hydrogels are often ultra soft (few hundreds of Pa) and certain chemical or physical agents are needed to increase the number of crosslinks and thus to raise gel’s stiffness. Glutaraldehyde, genipin and ribose are some of the compounds used to increase collagen gel’s strength^10–13^. Methacrylic anhydride along with a photoinitiator has also been employed as a crosslinker to modify collagen gel’s stiffness. The exposure of methacrylated collagen under UV light of a certain wavelength favors the linkage between methacrylated collagen fibers^14–16^. Thanks to the use of the above-mentioned techniques, collagen gels reaching a few kPa can be successfully made. The seeding of cells on top of compliant hydrogels, which present similar mechanical properties to cells, resembles more tissue-like environments; however, 3D dimensionality is still lacking^17,18^. For that reason, hydrogels made of some ECM proteins, such as collagen, are gaining importance to be used as scaffolds to culture cells^17,19–23^. Type I collagen is the most abundant protein in connective tissue that provides mechanical stability and strength and fibroblasts are responsible for its synthesis^24^; thus, its use resembling ECM properties is increasing.

Cells, in tissues, are surrounded by neighboring cells and ECM, which together provide different stresses and biochemical cues as in a 2D environment. Tissue availability is limited for research purposes, which is the closest biological organization to cells’ natural environment; therefore, other strategies are needed to provide tissue-like conditions. In disease, cells can experience changes in biochemical and biomechanical composition that may be transmitted to the ECM, procuring ECM composition and stiffness alterations as well^25^. Accordingly, both cells and ECM variations may generate feedback loop responses, in which cell-ECM interplay is essential. Hence, cell-to-cell interaction as well as with its surrounding is vital for cell mechanical properties development. Cell behavior assessment surrounded by its natural ECM is desirable; otherwise artificial hydrogels mimicking ECM composition and mechanics offer a good compromise^23,26–28^. 3D hydrogels provide physical constraints forcing the cells to adapt morphologically as well as chemically and biologically to an environment similar to that in tissue. There are several models to study fibroblasts contractile activity; free-floating matrix contraction and anchored matrix contraction are the most employed. In the method first the gel is detached from the substrate right away after gelation and is based on measuring the reduction in matrix’s diameter and the tension is distributed isotropically; however, the anchored model maintains the gel attach to the substrate for some time and after a certain time period, the gel is detached and measures changes in matrix height and the tension is distributed anisotropically. Many studies used the free-floating model because it is easier to measure; nevertheless, it was shown that fibroblasts do not proliferate in compliant matrices; they become arrested in cell phase G0^29,30^. Therefore, mechanical stress in the anchored matrix due to the attached surface seems to simulate better fibroblasts’ environment and more specifically wound healing process.

Cell and tissue mechanics have been investigated using several techniques, such as optical and magnetic tweezers, flow cytometer and atomic force microscopy (AFM)^31–35^. This latter technique has gained importance over the last twenty years, due to its ability to be used as a diagnostic tool as above mentioned. By applying an appropriate AFM methodology and contact model to analyze it, the sample’s mechanical response can be well described. AFM piezo-based nanopositioning provides the user flexibility to design any useful scheme to get the desired sample response. Step response^36^, force clamp^37^ and sweep modulation^38^ are some of the employed AFM methodologies to differentiate between elastic and viscous properties of cells. Fractional models, such as Kelvin-Voigt, and damping models based on exponential and power law’s cell response can be used to analyze AFM data^38–41^.

In this study fibroblasts from a patient suffering Dupuytren’s disease were used. Dupuytren’s disease is a fibromatosis of the connective tissue of the palm that generates nodules and cord in the palmar fascia leading to finger flexion and hand contraction^42–45^. Fibroblasts differentiation into myofibroblast phenotype (α-smooth muscle actin positive fibroblasts) is one of the disease’s characteristics that bring together changes in the ECM that enhances fibroblasts stress and thus hand contraction^46–49^. We dispose of three fibroblast types from different tissue samples from the palm of the same patient; healthy from dermal region, scar fibroblasts from the scar excision and Dupuytren fibroblasts from the nodules of the palmar fascia, presenting a myofibroblast phenotype^8^.

The aim of this project is to assess cells mechanical behavior with increasing substrate stiffness and cells contractile ability when embedded in collagen gels. For this purpose, firstly cells were seeded in collagen gels of different stiffness (0.2 to 2 kPa) and petri dishes as a hard substrate for comparison. Secondly, to provide 3D dimensionality, fibroblasts were embedded in soft collagen gels and gels’ stiffness were evaluated after different culture times.

## Results

Collagen from rat-tail was used to prepare collagen gels of two different stiffnesses. We used the same collagen concentration in both gels (5.5 mg/ml); however, to increase gel stiffness we employed a crosslinking molecule (methacrylic anhydride) to induce an increase in collagen fibers crosslinks. The free amino groups on the lysine residues of the collagen fibers undergo nucleophilic substitution with methacrylic anhydride. A photoinitiator, in our case Irgacure, when exposed to UV light (365 nm) generates radicals that are transmitted to methacrylate groups favoring the linkage between them in different collagen fibers increasing the number of crosslinks and therefore leading to stiffening of the gel (Fig. 1).

**Figure 1.**
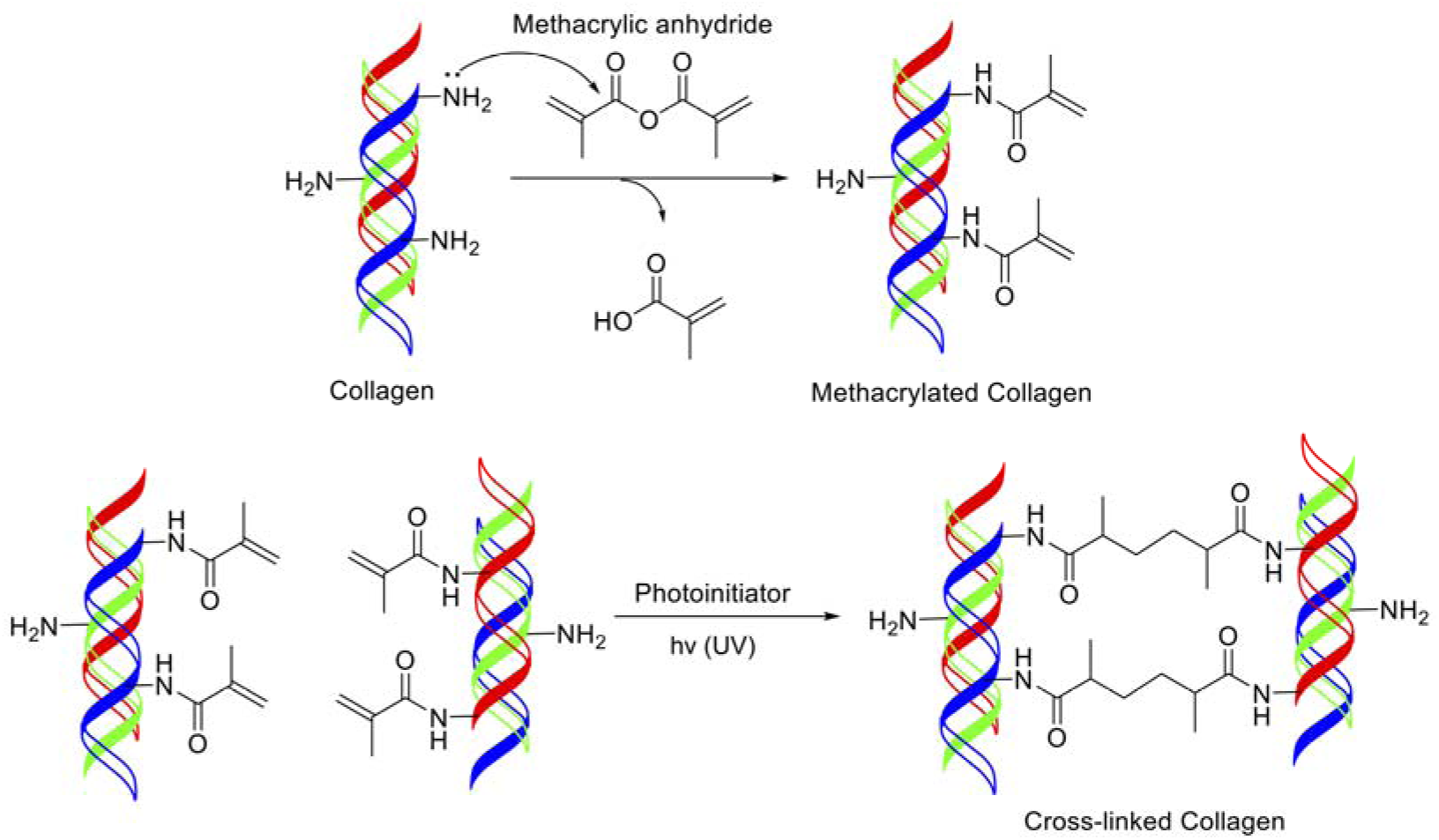
Synthesis of methacrylated collagen.

We performed AFM experiments to evaluate gel stiffness and we obtained an increase in stiffness of 10-fold of the methacrylated collagen with respect to the soft collagen (soft collagen: 0.2-0.3 kPa; methacrylated: 2-3 kPa) (Fig. 2).

**Figure 2.**
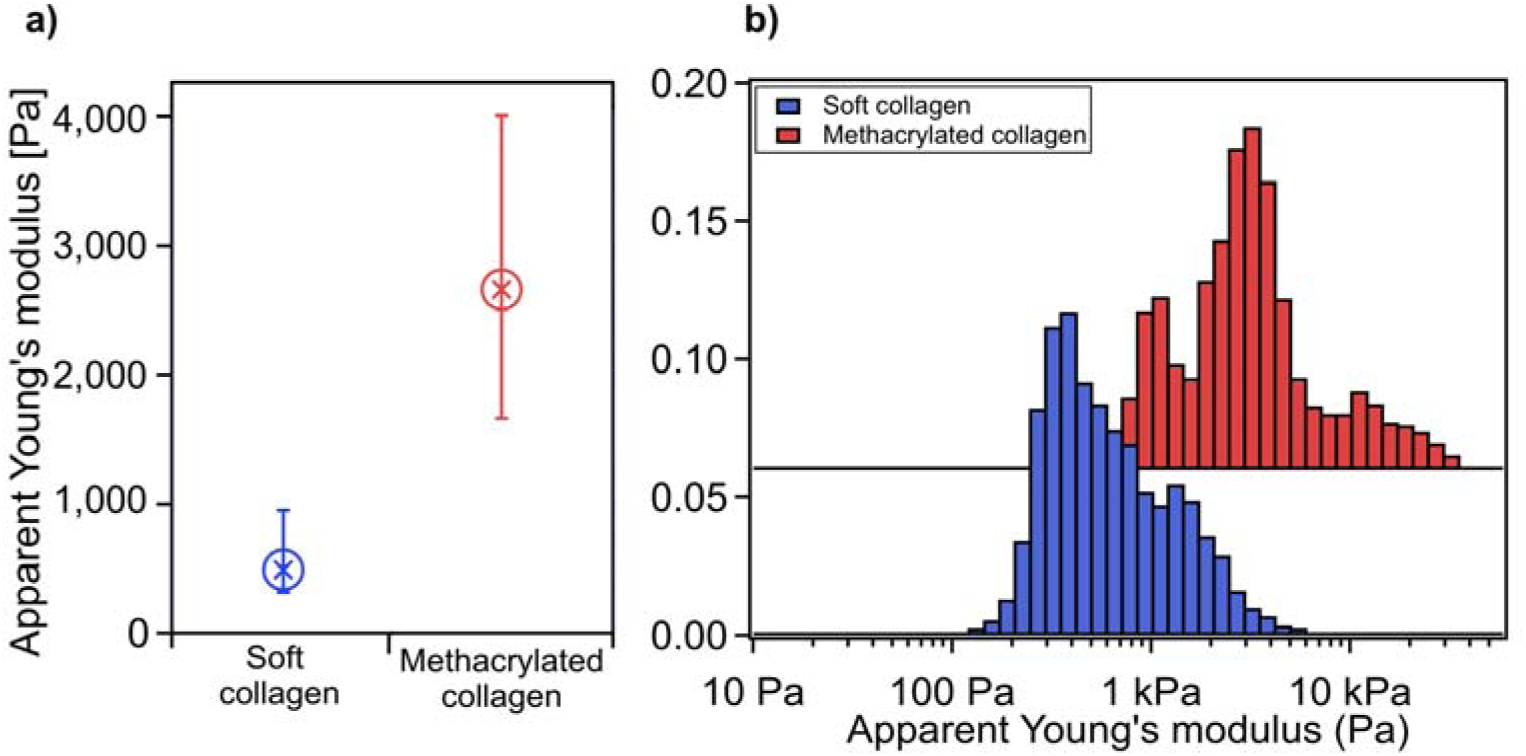
A) Box plot (median with 25/75 percentiles) of collagen gels stiffness (*n* = 7) and B) logarithmic histogram representation of apparent Young’s modulus. Data were extracted from AFM measurements on bare gels incubated the same time as cells.

We seeded three different fibroblast types on the two different collagen gels and used petri dishes as a hard substrate for comparison. Thanks to the collagen gels’ stiffness obtained and the petri dishes, we assessed cell mechanical behavior in substrates with really different stiffness, thus allowing us to encompass cell response in different environments. Cells data on hard petri dishes are used as a reference and were described in the previous work^50^.

As a comparative parameter, the apparent Young’s modulus from the approach curve was computed and the Hertz model for spherical indenters was used (Fig. 3). All cell types showed a decrease in apparent Young’s modulus when seeded in methacrylated collagen in comparison to soft collagen gels. Nevertheless, healthy and scar fibroblasts presented a slightly increase in apparent Young’s modulus when seeded on petri dishes with respect to methacrylated collagen, but Dupuytren fibroblasts showed a large increase in apparent Young’s modulus even surpassing apparent Young’s modulus values of the cells when seeded on soft collagen gels (Fig. 3 and Table 1). Statistical analysis showed significant differences among healthy fibroblasts seeded on the different substrates. Scar fibroblasts seeded on soft collagen and methacrylated collagen showed significant differences, as well as scar fibroblasts plated on soft collagen and petri dishes. Additionally, significant differences among Dupuytren fibroblasts seeded on the three different substrates were observed. Cohen’s d test was calculated to complement Wilcoxon test and large size effect was observed between healthy fibroblasts seeded on soft and methacrylated collagen as well as methacrylated and petri dish. Medium size effect was found between healthy fibroblasts seeded on soft collagen and petri dish. Moreover, medium size effect could be seen between scar fibroblasts seeded on soft collagen and petri dish, as well as when seeded on methacrylated collagen and petri dish. Scar fibroblasts seeded on soft and methacrylated collagen showed large size effect. Same size effect results as healthy fibroblasts were observed for Dupuytren fibroblasts, except when comparing Dupuytren fibroblasts seeded on soft and methacrylated collagen, in which medium size effect was obtained.

**Figure 3.**
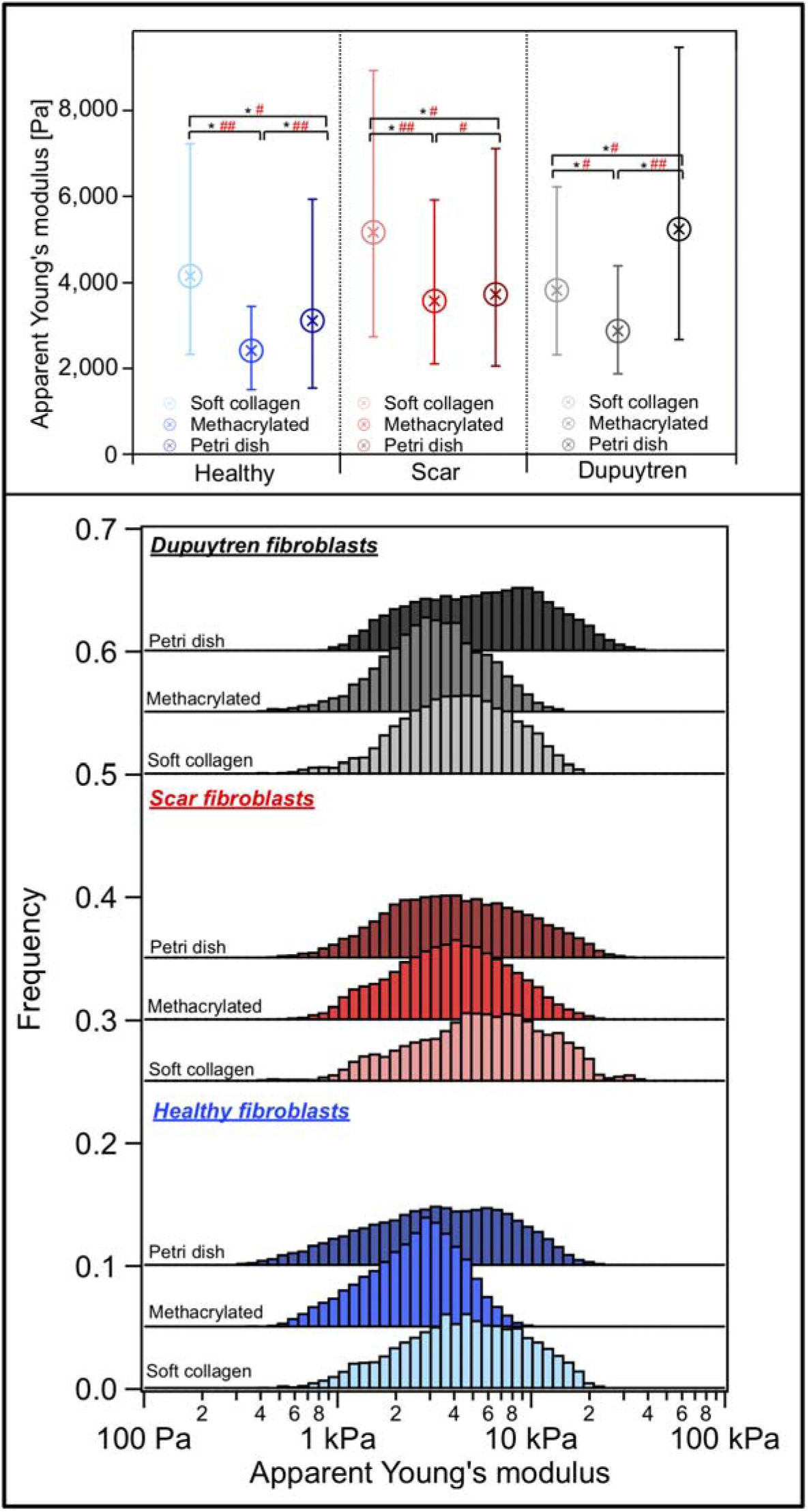
Box plot (median with 25/75 percentiles) and histogram distribution of the Apparent Young’s modulus from approach curves of the three different fibroblasts seeded on soft collagen gel, methacrylated collagen gel and petri dish (*n* = 30). Data are sorted by cell type and within each group; from lightest to darkest color represents the softest to the stiffest substrate where the cells were seeded.

**Table 1.** Numerical results of apparent Young’s modulus of fibroblasts, presented in Fig. 3 (median).

| **Apparent**<br>**Young's modulus** | **Soft collagen** | **Methacrylated**<br>**collagen** | **Petri dish** |
| --- | --- | --- | --- |
| **Healthy Fib.** | 4156 Pa | 2419 Pa | 3115 Pa |
| **Scar Fib.** | 5161 Pa | 3575 Pa | 3732 Pa |
| **Dupuytren Fib.** | 3826 Pa | 2876 Pa | 5236 Pa |

The hysteresis between approach and retract curves is due to cell viscosity, which cannot be quantified and separated from the elastic response in conventional force curves. To measure the elastic and viscous response of the cell samples, we used the sweep frequency scheme, previously described^52^. In figure S1 comparison between viscoelastic properties of the three different fibroblasts obtained from the sweep frequency data as a function of substrate were shown. In healthy and scar fibroblasts, the measured storage modulus at 1 Hz seemed independent on the substrate stiffness (healthy fibroblasts: 3150 Pa to 2500 Pa, from soft collagen to Petri dish; scar fibroblasts: 4050 Pa to 3050 Pa, from soft collagen to Petri dish), while Dupuytren fibroblasts showed an increase in storage modulus going from 3202 Pa (soft collagen) to 4322 Pa (Petri dish). The loss modulus at 1 Hz showed similar results as the storage modulus, in which healthy and scar fibroblasts slightly varied (healthy: 615 to 495 Pa, from soft collagen to Petri dish; and scar: 675 to 581 Pa, from soft collagen to Petri dish), whereas Dupuytren fibroblasts displayed an increase from 599 to 737 Pa (Fig. S1). Both moduli (storage and loss) displayed similar frequency dependence up to 10 Hz. However, loss modulus showed more marked frequency dependence at higher frequencies due to in a large extent the hydrodynamic drag of the cantilever in contact with the liquid (Fig. S2). Healthy and scar fibroblasts E* vs. frequency results almost overlapped each other, suggesting similar behavior regardless substrate stiffness. The representation of E* over frequency before and after viscous drag correction could be seen in figure S3. The three fibroblast types showed similar power law exponent in the different substrates, presenting values around 0.11, and loss tangent values were around 0.2 at 1 Hz (Fig. 4). E_0_ values varied depending on cell type and substrate and presented similar results as the storage modulus (Fig. S4) and the Newtonian viscous term (μ) was rather constant for Dupuytren fibroblasts regardless substrate stiffness (15 Pa·s) (Fig. S5).

**Figure 4.**
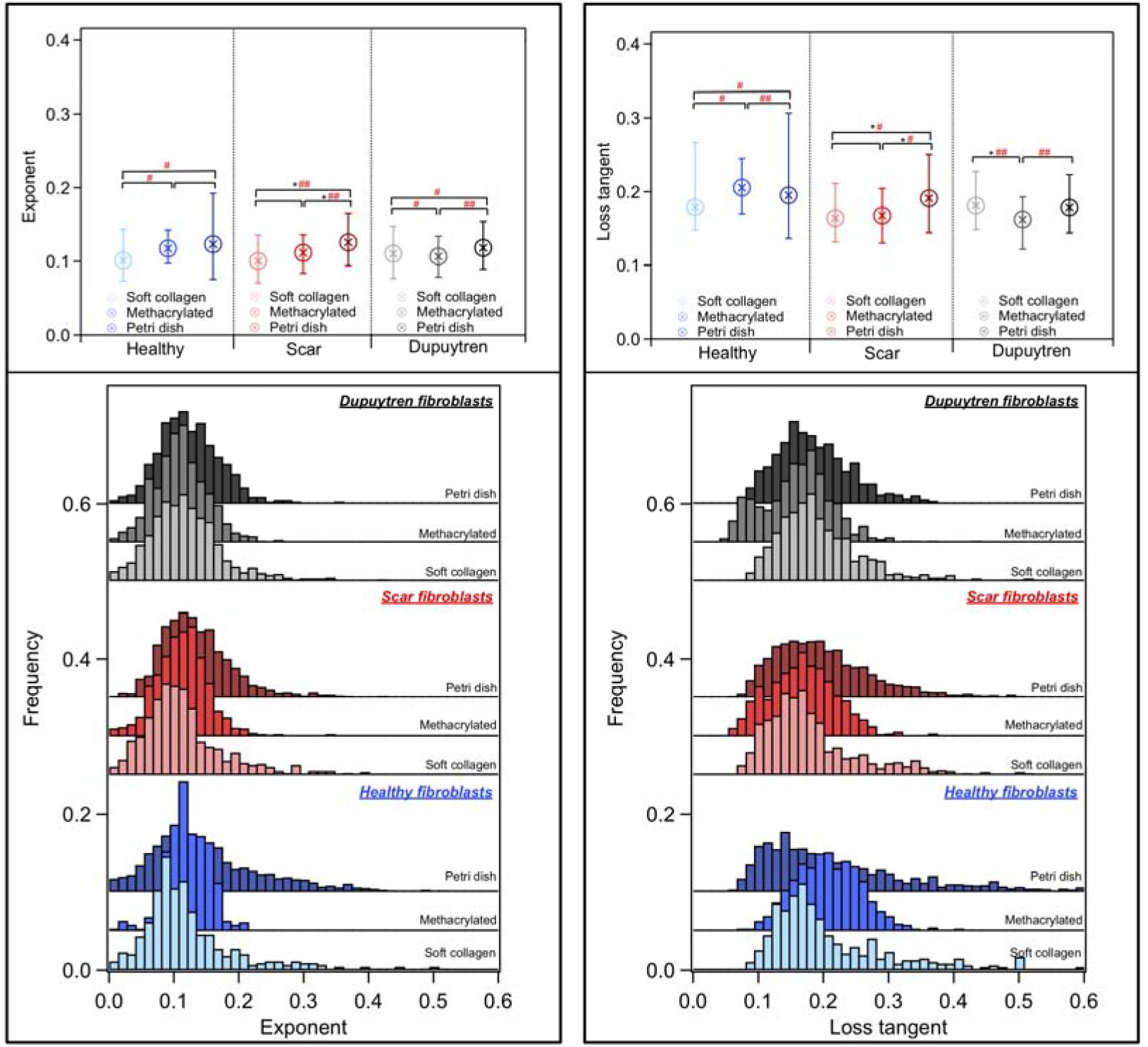
Box plot (median with 25/75 percentiles) and histogram distribution of the power law exponent and loss tangent at 1 Hz (left and right) (*n* = 30). Data are sorted by cell type and within each group; from lightest to darkest color represents the softest to the stiffest substrate where the cells were seeded.

Cytoskeleton organization in the cells seeded on the different substrates was evaluated labeling actin fibers. Scar and Dupuytren fibroblasts presented elongated shape and seemed to be aligned when seeded on soft collagen (Figure 5 d,g) while they displayed more spread body and randomly dispersed when seeded on methacrylated collagen (Figure 5 e,h) and petri dishes (Figure 5 f,i). Healthy fibroblasts seemed to follow the same trend; however, due to the low number of cells found in the images, big statements can not be made (Figure 5 a,b,c). Differences in cell morphology were also corroborated with the quantification of cell eccentricity and fibroblasts showed a decrease in eccentricity when seeded on petri dish with respect to soft collagen gel substrate (Fig. S6).

**Figure 5.**
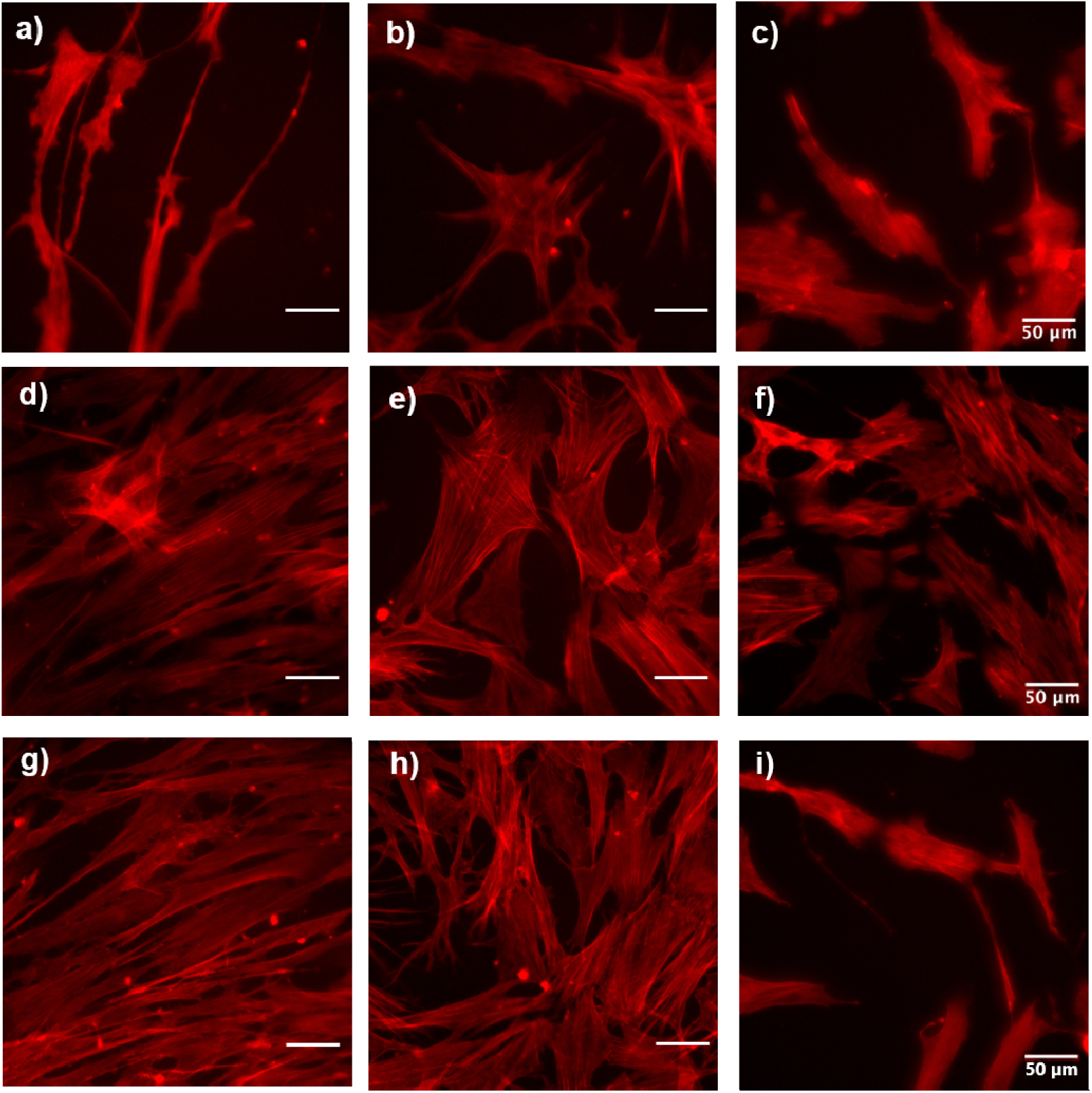
Fluorescence images of healthy, scar and Dupuytren fibroblasts labeled with rhodamine phalloidin (actin cytoskeleton). The upper panels show healthy fibroblasts seeded on soft collagen (a), methacrylated collagen (b) and Petri dish (c). The middle panels show scar fibroblasts, seeded on soft collagen (d), methacrylated collagen (e) and Petri dish (f). The lower panels show Dupuytren fibroblasts, seeded on soft collagen (g), methacrylated collagen (h) and Petri dish (i). Scale bars are 50 μm.

In an attempt to better resemble the cells’ environment in tissue, fibroblasts were embedded in soft collagen gels generating 3D structures. We measured 3D gels stiffness after three different incubation times: 2 days, 1 week and 2 weeks. Conventional force curves of the gels after the different incubation times were presented in figure 6. Note that there was an increase of the slope with increasing the incubation time in all gels regardless of fibroblasts type. This slope increment was observed in an increase in gel stiffness over time (Fig. 7).

**Figure 6.**
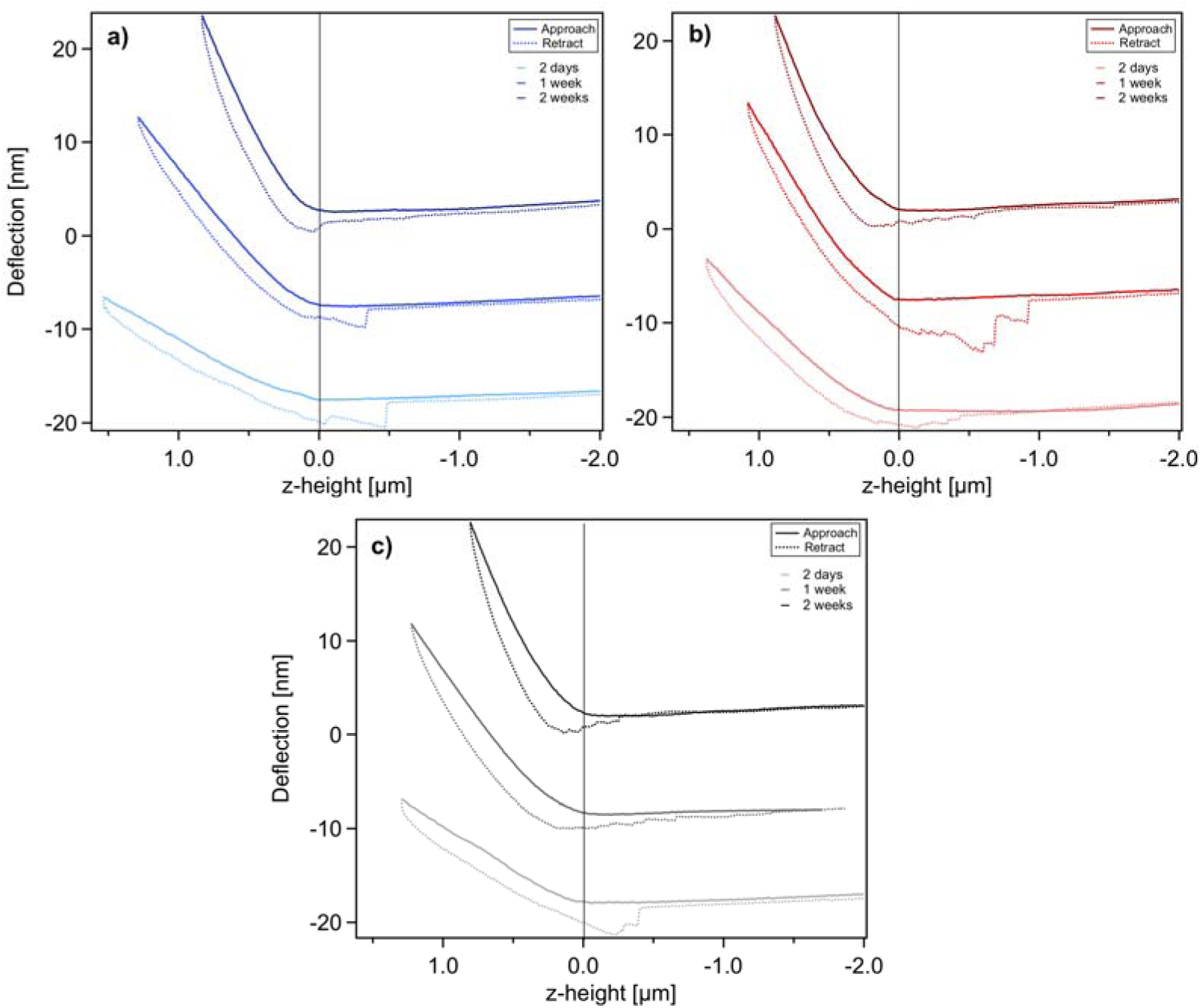
Representative force curves of gels where healthy (a), scar (b) and Dupuytren (c) were embedded. From lightest to darkest color in each graph represents gels measured after 2 days, 1 week and 2 weeks incubation time, respectively. Solid and dashed lines represent approach and retract curves, respectively.

**Figure 7.**
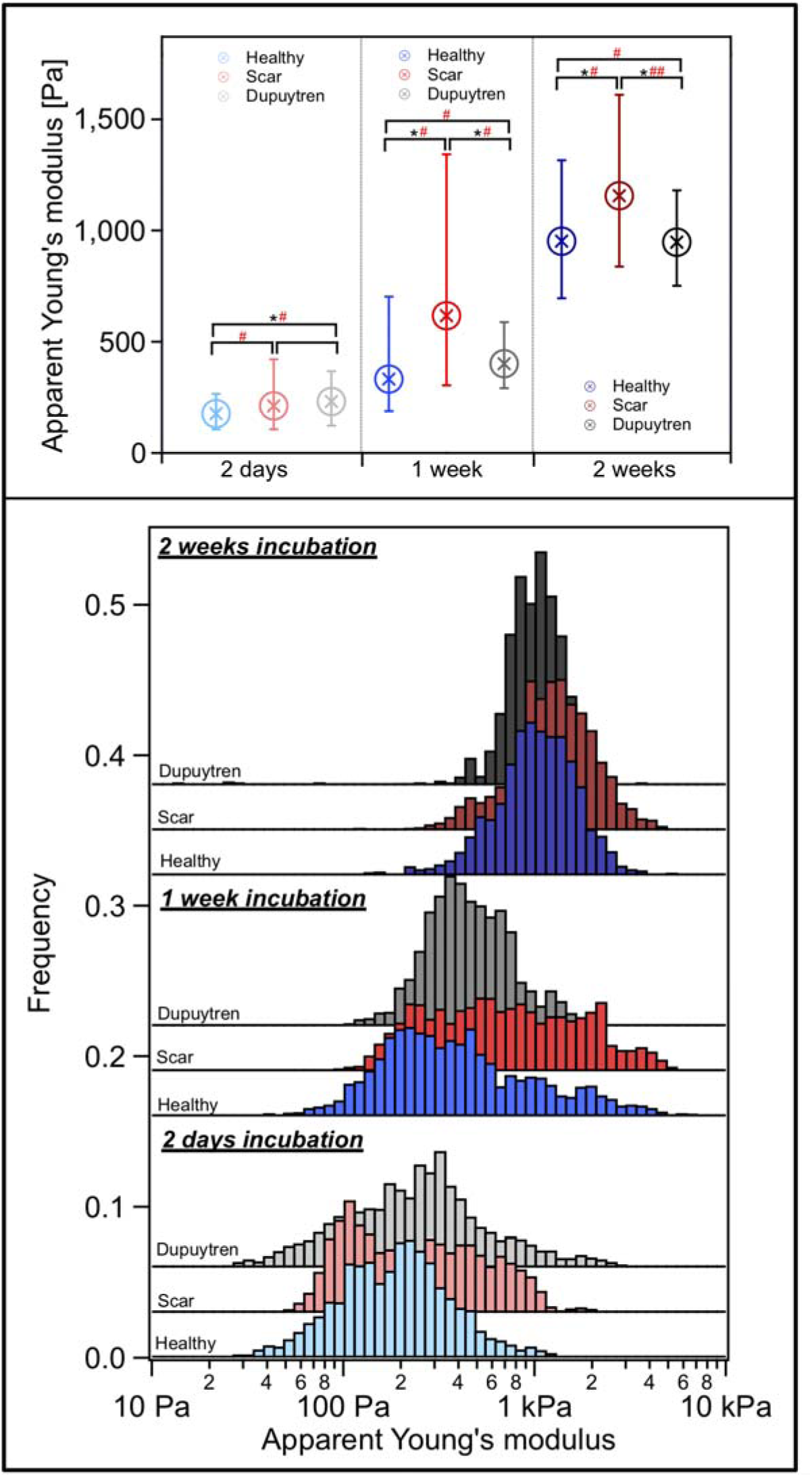
Box plot (median with 25/75 percentiles) and histogram distribution of the Apparent Young’s modulus from approach curves of gels (*n* = 30). Data sorted by incubation time. Within each group (incubation time) each color represents one type of fibroblast embedded in the gel (blue: healthy, red: scar and black: Dupuytren fibroblasts).

The measured apparent Young’s modulus of the gels went from 177 Pa (2 days) to 952 Pa (2 weeks) when healthy fibroblasts were embedded; 212 Pa (2 days) to 1156 Pa (2 weeks) for scar fibroblasts and 231 Pa (2 days) to 948 Pa (2 weeks) having Dupuytren fibroblasts embedded. Significant differences between healthy and Dupuytren gels after 2 days incubation could be seen in figure 7. In addition, healthy versus scar and scar versus Dupuytren gels after 1 week incubation also showed significant differences; these differences were also visible after 2 weeks of incubation. Cohen’s d analysis suggested medium size effect between healthy and scar; and healthy and Dupuytren gels after 2 days of incubation. A similar medium size effect could be observed among all gels after 1 week of incubation and healthy versus scar and healthy versus Dupuytren gels after 2 weeks of incubation. Finally, scar and Dupuytren gels after 2 weeks of incubation presented a large size effect.

We used sweep frequency data to obtain the viscoelastic response of the gels in the different conditions. Frequency dependence of storage and loss moduli are displayed in figure 8. We could see how storage and loss modulus almost follow the same trend over all frequency range after viscous drag correction (Fig. 8).

**Figure 8.**
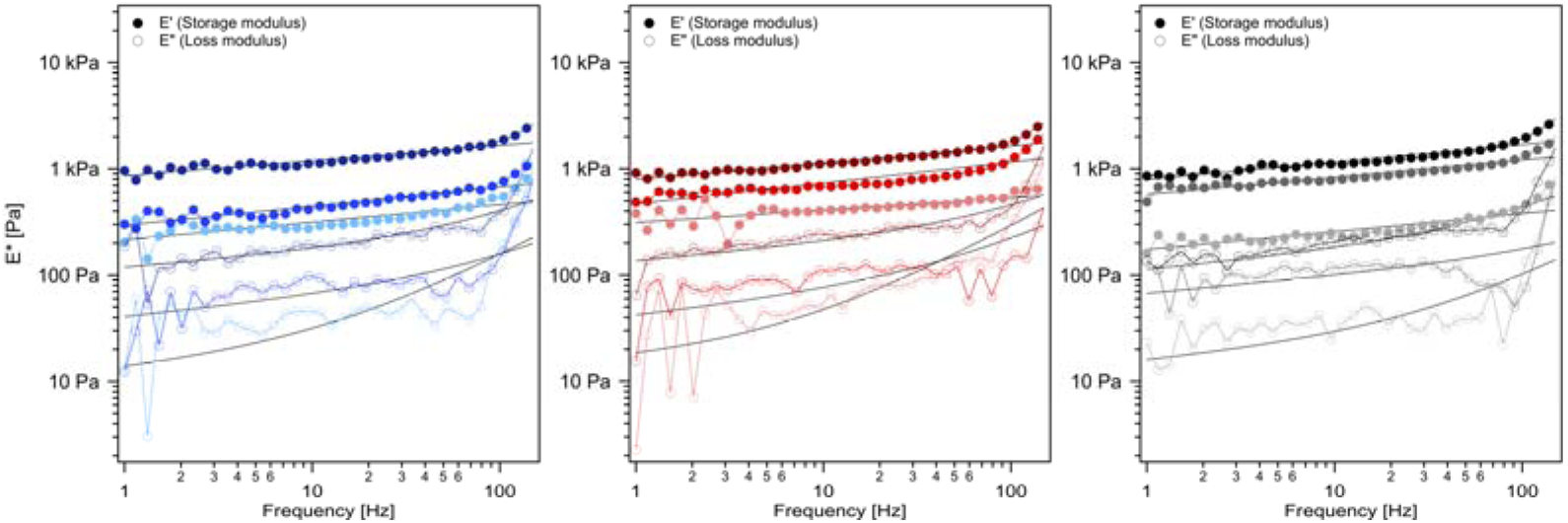
Frequency dependence of the storage modulus (filled symbols) and the loss modulus (open symbols) measured on gels in which healthy (blue), scar (red) and Dupuytren (black) fibroblasts (*n* = 30) were embedded at different oscillation frequencies (median). From the lightest to the darkest colors in each graph represents more to less incubation time (from 2 days to 2 weeks). Solid lines are the fit of the power-law structural damping model.

Storage modulus at 1 Hz versus power law exponent plot for all gels was presented in figure 9. We saw a tendency towards larger storage modulus values and slightly decrease in power law exponent, from 0.16 (2 days incubation) to 0.14 (2 weeks incubation) with increasing incubation time (Fig. S10).

**Figure 9.**
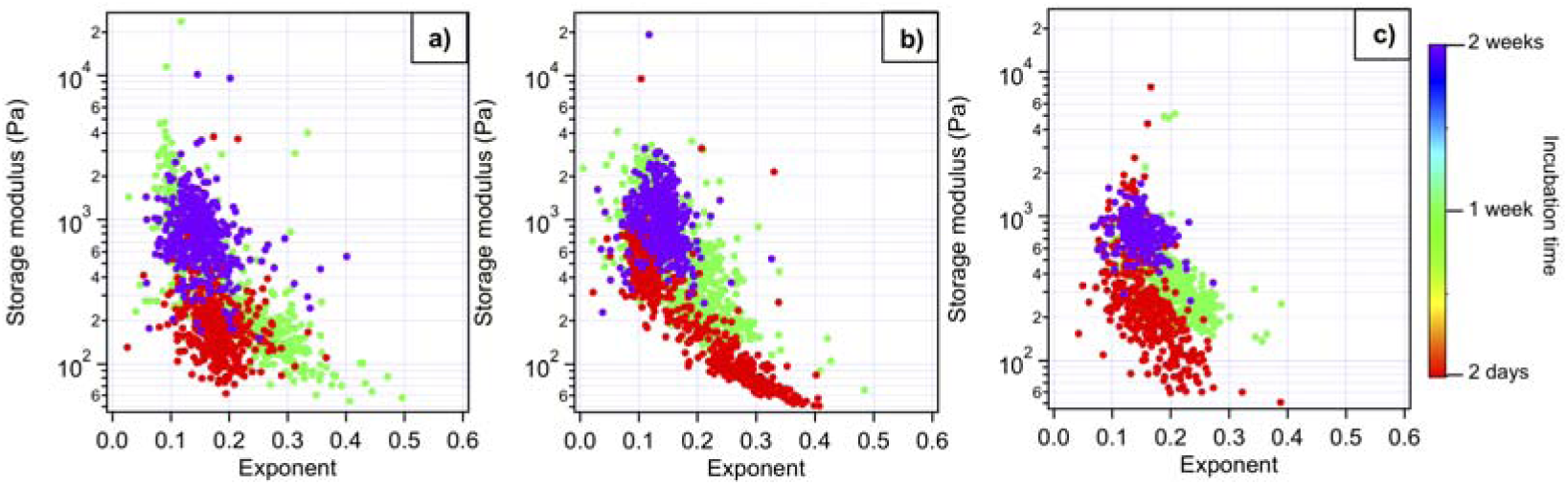
Scatterplot of storage modulus at 1 Hz versus power law exponent. Each dot corresponds to an individual force curve. a) Gels with healthy fibroblasts; b) gels with scar fibroblasts and c) gels with Dupuytren fibroblasts. Colors present gel data at different incubation times (*n* =30). Rheological properties of the gels can be seen in figures S7-S12.

There were no significant differences between gels in loss tangent parameter, presenting values close to 0.1. From less to more incubation time there was a small increase in loss tangent, related to an increase in gel viscosity (Fig. S9). E_0_ and Newtonian viscous term (μ) values were calculated and can be seen in figure S11-S12.

Live/dead staining of cells inside the collagen gels was performed in order to evaluate cell survival inside artificial hydrogels, showing the three fibroblasts a good viability when embedded in collagen gels (Fig. S13). Moreover, to assess how the cell’s cytoskeleton reacted to a soft 3D environment, actin fibers and nucleus were labeled (Fig. 10). We saw how Dupuytren fibroblasts spread faster than the other two cell types after 2 days of incubation (Figure 10 a,b,c); however, after 1 week and longer time periods, the three cell types seemed to stabilize, showing similar cell morphology and actin fibers distribution (Figure 10 d-i). Cell morphology was corroborated quantifying cell eccentricity, showing an increase in healthy and scar fibroblasts’ eccentricity over time but Dupuytren fibroblasts maintained similar values throughout the incubation period (Fig. S14).

**Figure 10.**
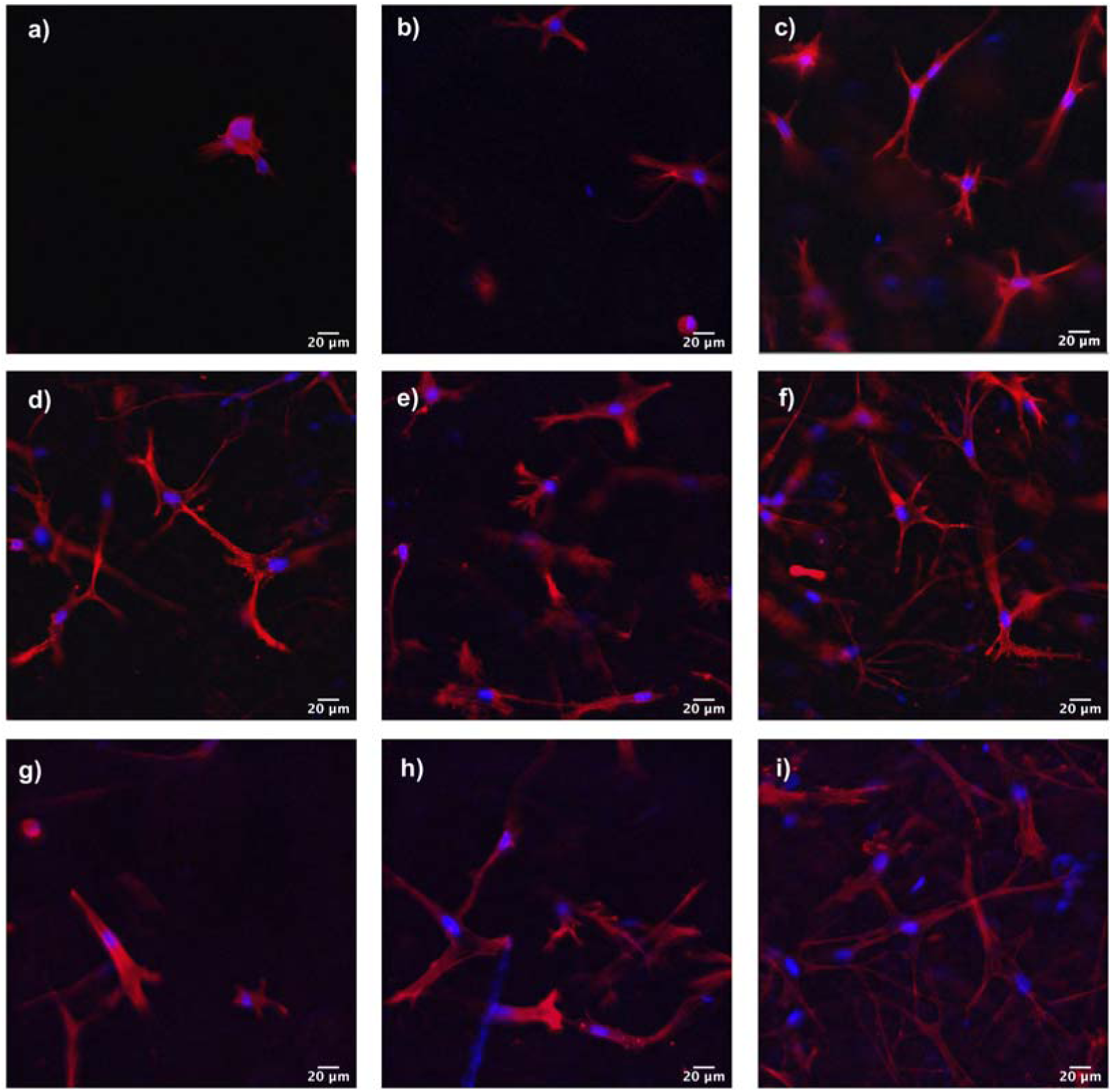
Fluorescence images of cells in 3D-collagen matrices. 2 days incubation: a) healthy, b) scar and c) Dupuytren fibroblasts; 1 week incubation: d) healthy, e) scar and f) Dupuytren fibroblasts and 2 weeks incubation: g) healthy, h) scar and i) Dupuytren fibroblasts. Actin fibers labeled in red and nucleus in blue. Scale bar: 20 µm.

We calculated the number of cells per cm^3^ inside each gel to verify how cells respond to the collagen 3D environment (Table 2). The number of healthy fibroblasts in the gel from 2 days to 1 week increased 3-fold and continued increasing constantly, 3-fold, from 1 week to 2 weeks incubation. Scar fibroblasts duplicated faster, increasing the number of cells 5-fold after 1 week. However, this rate decreased to 3-fold from 1 week to 2 weeks incubation. Dupuytren fibroblasts duplicated a bit faster than healthy from 2 days to 1 week (4-fold) and rose from 1 week to 2 weeks (4.5-fold).

**Table 2.**
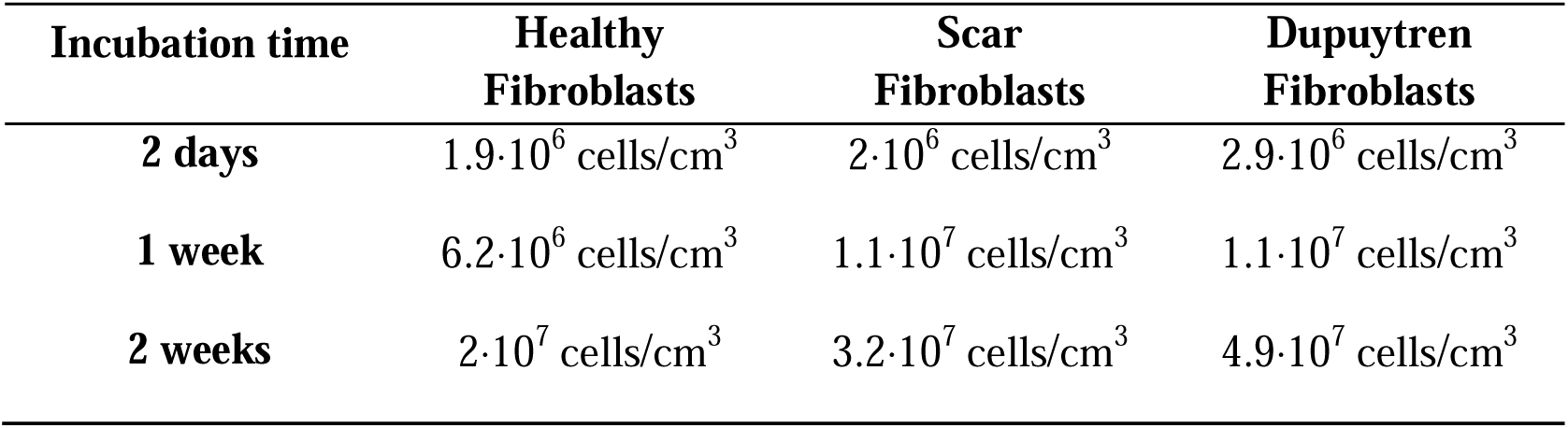
Number of cells per cm^3^ for each gel condition.

Immunostaining of the gels for fibronectin, collagen I and III was performed in combination to AFM measurements. Collagen I and III were not visible in any gel regardless of cell type and incubation time; however, differences in fibronectin secretion were observed. All cell types increased the secretion of fibronectin over time, presenting 2 weeks incubation the strongest signal (Fig. 11). 2 weeks incubation gels, in which Dupuytren fibroblasts were embedded, presented a slightly increase in fibronectin signal intensity. Besides, fibronectin signal was stronger on the upper layer of the gels, probably due to the impediments of the antibodies to diffuse within the gel.

**Figure 11.**
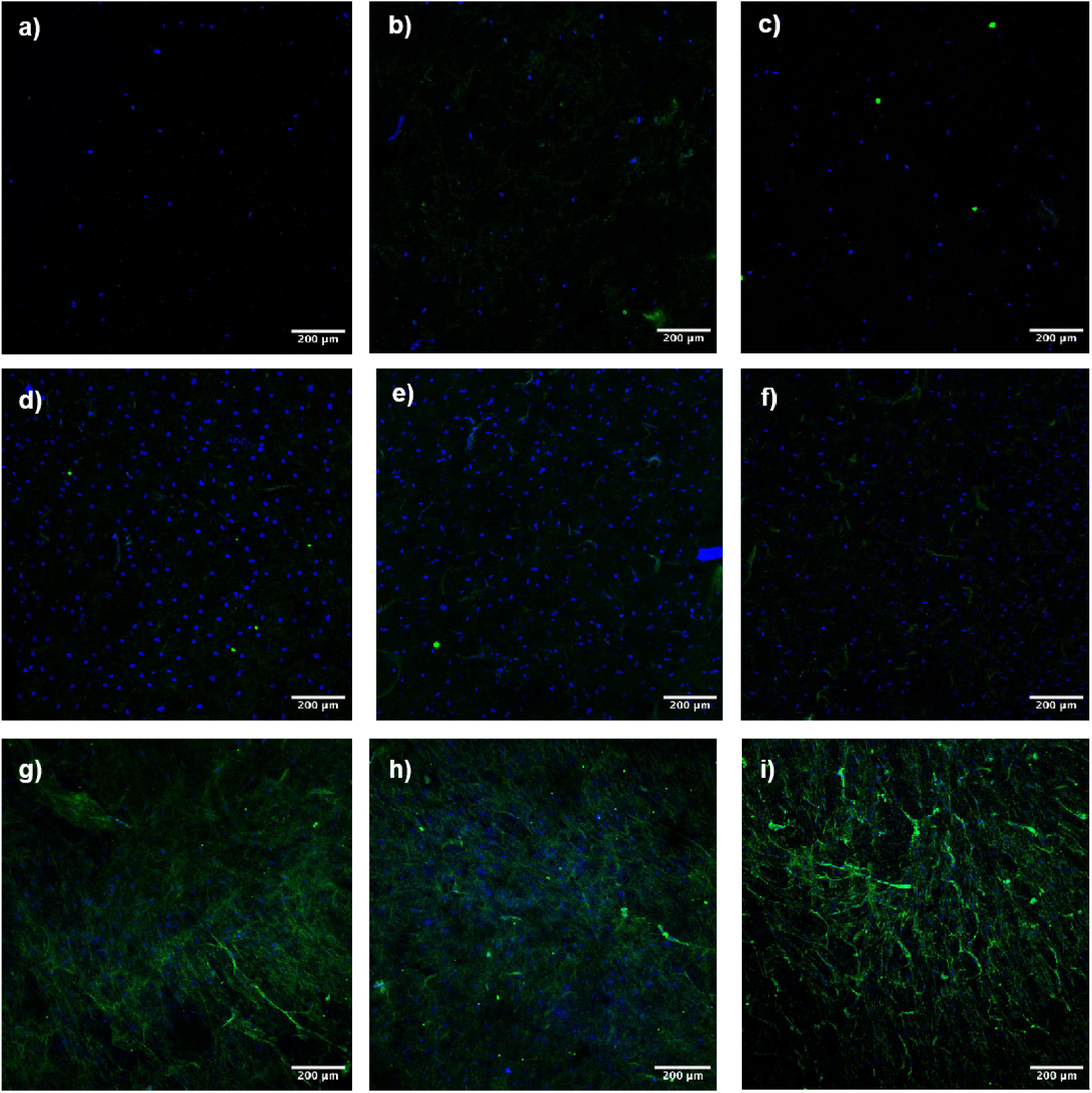
Fluorescence images of the 3D-collagen gels presenting fibronectin labeling in green and cell’s nucleus in blue (scale bar: 200 µm). 2 days incubation: a) healthy, b) scar and c) Dupuytren fibroblasts; 1 week incubation: d) healthy, e) scar and f) Dupuytren fibroblasts and 2 weeks incubation: g) healthy, h) scar and i) Dupuytren fibroblasts.

## Discussion

Collagen is the most abundant structural protein in the ECM and it is found in connective tissue such as cartilage, tendons, ligaments and skin. Fibroblasts are responsible of producing and remodeling collagen that is secreted to the ECM via exocytosis. It has been observed that artificial collagen gels use to be much softer than tissues or cells, and it may be due to lower amount of crosslinking between filaments in these artificial gels and the absence of a mixture of proteins that provide distinct strength. Different approaches were used to increase collagen gels stiffness - apart from increasing the concentration - and the most employed one is to use a photo-crosslinker that increment the number of crosslinks between collagen fibers, leading to a certain increase in gel stiffness. We used methacrylic anhydride and Irgacure and reached a 10-fold increase in apparent Young’s modulus with respect to the untreated collagen gel (soft collagen). How ECM mechanics and biochemical composition influences the cell’s mechanics is an important issue; therefore, the substrates used in this work can span a broad fibroblasts response due the different mechanical and biochemical composition.

In our experiments, we varied two experimental parameters: cell type and substrate stiffness, which resulted in a highly complex and interesting research project; therefore, it is possible to discuss it in two different ways. By focusing on individual fibroblast types seeded in different substrates, we saw that healthy and scar fibroblasts presented a similar response when changing the stiffness of the underlying support; showing slightly higher apparent Young’s modulus when seeded on soft collagen but they stayed almost unaltered when cultured in the methacrylated collagen and petri dish. These results may suggest that healthy and scar fibroblast present a similar phenotype and they react similarly to changes in substrate stiffness. This is an interesting finding, since it is unclear if scar fibroblasts resemble more healthy-like or pathological-like fibroblasts’s behavior. However, although Dupuytren fibroblasts also presented larger apparent Young’s modulus when seeded on soft collagen gels with respect to methacrylated collagen gels, its mechanical properties in hard petri dishes surpassed the rest, which corroborate their myofibroblast phenotype. Myofibroblasts are contractile fibroblasts developed in wound healing and facilitate wound closure^53–55^; thus, Dupuytren fibroblasts’ response to substrate stiffness is linked to the wound healing process. It may be noted that soft and methacrylated collagen gels present not only differences in stiffness but also in biochemical composition, therefore, changes in fibroblasts’ mechanics may not only depend on the stiffness of the underlying support but also on the biochemical composition. In fact, all fibroblasts displayed a decrease in stiffness when plated on methacrylated collagen, which may be due to the toxicity of methacrylate groups, to which cells were exposed. Previously, physiological matrices with different protein content as well as different stiffness were used to study fibroblasts response^56^. Pathological fibroblasts remodeled the ECM, leading to a stiffening of the matrix but healthy fibroblasts softened it. The matrices employed in the latter work presented more than one ECM protein; however, in our case, collagen I was the only ECM protein employed. We believe that our experimental setup provides clearer results regarding cell response to ECM stiffness because only one parameter is being investigated at the same time, which is ECM stiffness and the biochemical composition is not different enough to play a key role in our experiments. By looking at cells’ responses to the same substrate, curious results were obtained. In both collagen gels, fibroblasts followed the same distribution, in which all fibroblasts presented close values in terms of apparent Young’s modulus. Nonetheless, when cells were plated on hard petri dishes, an increased tendency towards larger apparent Young’s modulus values from healthy to Dupuytren fibroblasts were observed. The mechanical stress of a matrix is transmitted to the cell through integrins that connect the ECM to the cell actomyosin network via focal adhesion complexes^57–59^. Myofibroblasts present mature focal adhesion complexes that connect the cell cytoskeleton to the ECM and are essential for cell attachment; thus, they are more sensible to changes in ECM stiffness. The tendency towards larger apparent Young’s modulus values in fibroblasts seeded in hard petri dishes is related to the myofibroblast phenotype expressed by Dupuytren fibroblasts that adapt to variations in the mechanical properties of the surrounded environment. These results are consistent with previous findings, in which Dupuytren fibroblasts modified their mechanics according to the stiffness of PA gels^8^.

Frequency dependence of storage and loss modulus of all fibroblasts in all substrates present the same distribution, storage and loss modulus showing the same frequency dependence up to ≈ 10 Hz; however, loss modulus present a more marked frequency dependence at higher frequencies. Hydrodynamic viscous drag was corrected; therefore the more marked increase in loss modulus over frequency is an intrinsic behavior of cells. Cell viscosity is governed by internal friction of micro- and macromolecules sliding in the cytosol. Molecules of different dimensions contribute differently to the viscous response, being largest molecules the ones that contribute most^60^. Surprisingly, power law exponent and loss tangent values were constant across all cells and substrates. Fibroblasts solid-like behavior does not vary depending on the substrate stiffness, displaying close loss tangent values on soft collagen gels and hard petri dishes. These values corroborate the altered cell behavior in soft collagen gels that is similar to that in hard petri dishes, suggesting that substrate stiffness is not the only parameter governing cell mechanics, but that other hidden mechanisms are probably at play.

It is often possible to correlate cell mechanics to cell morphological changes; e.g. cells presenting elongated and spread shapes usually display higher apparent Young’s modulus that is related to the cytoskeleton development. It was known that cells increase their spreading with increasing the stiffness of the support, expecting to see higher cell spreading in petri dish substrates^61^. Cells plated in soft ECM functionalized PA gels (few hundreds of Pa), remain little spread with a rounded shape^62,63^. The three fibroblasts investigated in this work presented similar morphology and cytoskeleton organization in the different substrates regardless substrate stiffness. Although cells displayed a more orientated/aligned shape on soft collagen gels, cell spreading did not differ substantially among the different substrates. Cell eccentricity was calculated and surprisingly, the larger values were displayed by fibroblasts seeded on soft collagen gels. This cell behavior on soft collagen gels was also shown in 3T3 and MKF fibroblasts that presented similar morphological properties in compliant collagen gels and rigid glass substrates^64^.

Cell’s natural environment provides of 3D dimensionality as they are wrapped in ECM that restricts cell mobility and spreading, and causing a downstream signaling pathway within the cells that activates biochemical and biophysical cues that may lead to a cell response and therefore changes in cell mechanics. Changes in cell mechanics are also transmitted to the ECM, generating ECM release/contraction or changes in ECM composition and/or organization. To provide the cells of these 3D environmental stimuli, fibroblasts were embedded in soft collagen gels. Gels were anchored to coverslips since gelation was performed directly on them, thus subjecting them to an additional tension. Our experimental setup resembles to the anchored matrix contraction model, in which 3D collagen gels are attached to a surface for up to a certain time period allowing possible myofibroblast generation and then the gel is released from the surface. This model is needed when myofibroblast differentiation wants to be study. The free-floating model does not provide sufficient tension to stimulate myofibroblast differentiation; thus, it is really important to choose the appropriate model for the desired results. Matrix dimensional changes to assess fibroblasts tension generation have been studied using light microscopy or optical coherence tomography (OCT) and anchored collagen matrix model^20^. In our experimental work, changes in gels mechanical properties were assessed using the AFM. AFM allows us to obtain a quantitative measure of gel’s tensional variations rather than changes in matrix’s dimensions. Gel contraction occurs as a consequence of motile activity by cells trying to migrate through the matrix, termed tractional remodeling. Moreover, gel contraction also occurs as fibroblasts spread, duplicate and elongate, as they first organize proximal collagen fibrils and subsequently these tractional forces are propagated throughout the entire collagen fibril network, resulting in the whole matrix contraction^20,22,48^. Changes in gel stiffness over time were expected; however, surprisingly, there were no big differences in apparent Young’s modulus among gels populated by the different fibroblasts. We may see a small difference in apparent Young’s modulus between healthy and Dupuytren gels (being the latest slightly stiffer) after 2 days of incubation. This is understandable since Dupuytren fibroblasts present more spread morphology than healthy fibroblasts and cell density is larger by a million cells, increasing cell-ECM contraction. However, after 2 weeks of incubation, gels’ stiffness stabilizes, displaying no differences among the different fibroblasts.

We would have expected to see differences between gel stiffness when different types of fibroblasts were inserted; however, that is not the case in our results. A possible explanation may be that the number of cells inserted was not enough to produce a readable gel contraction and change in stiffness or the stiffness of the gel itself was low to activate cell’s response. Besides, it could also be that the technology employed to characterize it (AFM) was not the most appropriate. AFM technique allows the use of cantilevers of different geometries and dimensions. We employed a spherical tip with a 5.5 µm radius, considering it a good compromise between spatial resolution and large scan size. Moreover, a large tip radius, like the one we use, needs to apply large forces to reach reasonable indentation depths. Even then, since our hydrogels have around 250 µm thickness, the AFM tip only records surface responses, which may not be sufficient to get a good value for the general gel stiffness.

Rheological properties of the gels were measured and the frequency dependence of the storage and loss modulus showed almost identical distribution. Loss modulus is almost 10-fold smaller than the storage modulus, indicating that these gels have properties close to pure elastic materials due to the low viscous contribution they have in comparison to cells. After 10 Hz, loss modulus distribution does not show same frequency dependence as cells. The more marked frequency dependence of loss modulus shown by cells is linked to the intrinsic viscous behavior of cells related to the presence of micro and macromolecules that cause friction, like proteins and organelles in the cell’s cytosol that actually is really different to the composition of the artificial collagen hydrogels. The larger the molecule, the larger the viscous contribution^60^. Loss tangent values are similar in all gels at the same incubation time but there is a slight increase over time, related to an increase in the liquid-like behavior, which can be related to an increase in cell number and/or spreading. This small change in gel’s loss tangent towards more liquid-like behavior resembles cell interior, in which embedded cells mimic the micro and macromolecules in the cytosol. An increment in cell number, provides an increase in frictional elements, thus in viscosity. Nevertheless, loss tangent values of the gels are still smaller than the cell’s values, indicating a more elastic/solid-like behavior. Similar loss tangent values were observed in PDMS gels^65^. Storage modulus’ increase of the gels over time has been seen with a rather constant power law exponent. The almost constant values of power law exponent of the gels over time suggests that gel’s rheological changes after two weeks of incubation may not be sufficient to procure substantial changes that are readable with our equipment.

The large difference in cell number inside each gel does not procure substantial mechanical changes. At the longest incubation time, apart from an increase in cell number inside the gels, an increase in cell deposition on the upper layer of the gel, presenting elongated morphologies similar to the 2D images, was observed. The accumulation of cells on the top of the gels may suggest a cells’ tendency not to stay inside the collagen gel, migrating to the upper layer. It may appear that the fibroblast’s nature leads to leaving the resting state they acquired inside the 3D gel towards a more contractile behavior in the 2D environment, behaving similarly to wound healing process.

As above mentioned, healthy and scar fibroblasts present more rounded shape than Dupuytren fibroblasts at short incubation times (2 days). Nonetheless, at longer incubation times all fibroblasts display similar morphological features showing membrane protrusions, resembling dendritic extensions^66^. Fibroblasts exhibiting dendritic/bipolar morphologies resemble fibroblasts in tissue under resting conditions^67–71^ while cells with well-defined stress fibers are typically observed in 2D conditions, like in petri dishes or during activated conditions such as wound repair and fibrosis^72–74^. Those differences in cell dendritic protrusions and well-defined stress fibers can be observed in figure 6 and figure 11, representing fibroblasts in 2D and 3D collagen matrices, respectively. Furthermore, it was observed that fibroblasts stress is linked to the stiffness of the matrix, needing a certain matrix stiffness to develop stress fibers and later on differentiate into myofibroblast phenotype^55^. Our fibroblasts morphological features in soft 3D collagen gels suggests that they present a resting state as the stiffness of the gel is not enough to stimulate fibroblast activation and therefore myofibroblast differentiation. Moreover, our gels’ mechanical properties (too soft) do not allow distinguishing between healthy and “pathological” Dupuytren fibroblasts’ contractile properties either.

AFM measurements were combined with immunostaining for fibronectin. In natural conditions, fibroblasts secrete fibronectin along with collagen I and III. The absence of collagen I and collagen III in our data, even after 2 weeks of incubation, may suggest that the formation and secretion of those proteins by fibroblasts is produced on longer time scales. Different incubation times for both primary and secondary antibodies were applied, finally improving fibronectin signals using longer incubation time periods. Protocols similar to whole mount staining, which is used for thick tissue samples such as embryos, did not provide better results, as well as sirius red staining (employed to stain and distinguish collagen from the rest of the sample components). Fibroblasts are one of the most important cells that secrete fibronectin to the matrix and it is a key factor for wound healing. Fibronectin helps in the formation of a proper matrix for cell’s migration and growth during the development of granulation tissue, connective tissue. Besides collagen deposition at the wounded area is developed with the help of fibronectin. Therefore, fibronectin secretion by fibroblasts before collagen deposition is in agreement with our results. We saw how fibronectin signal increases over incubation time, from no signal after 2 days of incubation to visible fibers after 2 weeks of incubation. Moreover, gels in which Dupuytren fibroblasts were embedded seem to present clearer fibronectin filaments over the other fibroblasts. This corroborates the pathological myofibroblast phenotype of Dupuytren fibroblasts that secrete more fibronectin to the matrix than the other two cell types. Changes in fibronectin expression were also observed in cancer and its increment was seen in lung carcinoma^75^.

## Conclusion

The modulation of the mechanical properties of collagen gels was possible thanks to the addition of methacrylic anhydride, a crosslinker that increases the number of crosslinks between collagen fibers leading to an increase of the gel’s stiffness. Healthy and scar fibroblasts behaved similarly when changing the mechanical properties of the underlying support, suggesting that the scar fibroblasts’ mechanical properties under these specific conditions resembled more those of healthy fibroblasts. Moreover, the apparent Young’s modulus of these fibroblasts when seeded on soft collagen gels was similar or even higher than when seeded on hard Petri dishes. However, Dupuytren fibroblasts modified its stiffness to that of the substrate, supporting its myofibroblast phenotype. Furthermore, the morphology of fibroblasts in soft collagen gels closely resembled that in hard Petri dishes, suggesting that the morphological and mechanical properties of our fibroblasts are not only influenced by the substrate’s stiffness but also by the biochemical composition. The different fibroblast types embedded in soft 3D-collagen gels did not produce large differences in gel mechanical properties; thus, gel tension. Dupuytren fibroblasts spread faster at shorter time periods; nevertheless, at longer time periods all fibroblasts presented similar morphology and contraction properties. The stiffness of the 3D collagen gels prepared was not sufficient to activate the contractile properties of fibroblasts; hence, cells did not differentiate to myofibroblasts; thus, the fibroblasts presented dendritic protrusions as if they were in resting state. This study corroborates the fact that myofibroblast differentiation is governed by changes in the stiffness of the surrounding environment and is facilitated by stiffer environments. Our study encompasses a wide range of cell response, from 2D to 3D environment, increasing the resemblance of tissue-like conditions, which is closer to the cell’s natural environment. We believe that artificial hydrogels made of some representative ECM protein, such as collagen, are a good compromise to study cell behavior in 3D.

## Materials and Methods

### Collagen extraction

Tails of Sprague-Dawley rats (male, 350 g) were collected as a by-product from other experiments in the animal facilities of the School of Medicine (University of Barcelona) and tendons were extracted to obtain a type I collagen solution by following the protocol by Rajan and coworkers^76^. Briefly, tendons were washed with 100 % acetone and 70 % isopropanol for five minutes each. Washed tendons were then dissolved in 0.02 M acetic acid and kept for 48-72 hours at 4°C under slow stirring until complete dissolution. Collagen fibers were triturated and kept frozen at −80°C. Finally, frozen collagen was lyophilized and kept at −80°C until used. 5.5 mg/ml collagen solution in 0.02 M acetic acid was employed.

### Gel preparation

Collagen I from rat-tail was used to generate collagen gels of different stiffness. We called “soft” to collagen gels with a concentration of 5.5 mg/ml in 0.02 M acetic acid and “methacrylated” to the methacrylated collagen. Methacrylated collagen increases its stiffness with respect to the soft collagen up to 10%. Briefly, 5.5 mg/ml collagen was mixed with 1/9 of PBS 10x and 19 µL NaOH 1M. The solution was mixed and 6 % of methacrylic anhydride (Sigma), 9 µL of Irgacure initiator (Advanced Biomatrix) and 19 µL NaOH 1M was added. The solution was vortexed and a neutral pH around 7-8 should be obtained. Collagen solution (either soft collagen or methacrylated) was poured in a hydrophilic 24 mm square coverslip and covered with a hydrophobic 22 mm ø circular coverslip to generate a flat surface and to get a homogeneous gel height. The collagen solution between the coverslips was incubated for one hour at 37°C at 5 % CO_2_ for polymerization and once the gel was formed; the top coverslip can be removed adding PBS to detach it easily. Methacrylated gels then were exposed to 10 minutes UV light to increase gel stiffness.

3D-gels with cells in it were prepared as follows: first, collagen gel solution was made: 5.5 mg/ml collagen was mixed with 1/9 of PBS 10x and 19 µL NaOH 1M. The solution was vortexed and a neutral pH around 7-8 should be obtained. Then 4·10^5^ cells/ml were prepared and added to the collagen solution. 460 µL of collagen solution were mixed with 40 µL of cells solution. The final cells’ concentration used was 8·10^5^ cells/cm^3^. Then, 250 µL of the mixed solution were poured in the 24 mm square coverslips and covered with the 22 mm ø coverslips to generate a homogenous flat surface for AFM experiments. The mixture was incubated for 1 hour at 37°C and 5 % CO_2_. Finally, the top coverslip was released and the gels were immersed in cell culture medium and incubated until AFM experiments were performed.

### Coverslips functionalization

24 mm square coverslips were functionalized to make them positively charged. Coverslips were washed with 100 % ethanol and exposed to ambient air until dried. Then, they were incubated with NaOH 0.1M and (N-[3(Trimethoxysilyl)propyl]ethylenediamine) (Sigma Merck) for 3 minutes each step. Coverslips were then washed three times with milliQ water for 10 minutes each step and incubated with 0.5 % glutaraldehyde for 30 minutes. Finally, they were washed three times with milliQ water for 10 minutes each step and exposed to ambient air until they were completely dried.

22 mm ø circular coverslips were functionalized to make them hydrophobic. Coverslips were washed with 100 % ethanol and dried. Then, they were covered with sigmacote solution (Sigma) for 1 minute and washed three times with milliQ water for 10 minutes each step. Finally, the coverslips were exposed to ambient air to dry.

### Cell culture

Three types of primary fibroblasts from the palm of the same patient who suffered from Dupuytren’s disease were used in this work: healthy fibroblasts from the dermis of the palm, scar fibroblasts from a wounded area, and Dupuytren fibroblasts from the palmar fascia. All cell types were cultured in DMEM medium (containing 4.5 g/L D-glucose, FG0435, Sigma) and incubated at 37°C in a humidified atmosphere of 95 % air and 5 % CO2. Medium was supplemented with 10 % FBS (fetal bovine serum) (F7524, Sigma) and 2 % penicillin-streptomycin (P0781, Sigma). Prior cell seeding, 2D-collagen gels were incubated with medium for 30 minutes to promote serum protein absorption on the gels, hence, cell adhesion. Cells were seeded 48 hours prior to the AFM measurements. 3D-collagen gels were incubated 2 days, 1 week or 2 weeks prior to AFM experiments. Fibroblasts between passages 8-12 were used for all the experiments. The study was conducted in accordance with the Declaration of Helsinki, and approved by the local Ethics Committee (ACrztekammer Bremen, #336/2012). Patient was informed pre-operatively and had given its informed consent to anonymous tissue donation.

### AFM experiments

AFM experiments were performed with a MFP3D AFM (Asylum Research, Santa Barbara, CA, USA) to measure mechanical properties of fibroblasts in 2D- and 3D-environment. An optical microscope was combined with the AFM to be able to control tips and samples (Zeiss Axiovert 135, Zeiss, Oberkochen). PFQNM cantilevers presenting three-sided pyramidal tips with 65-75 nm tip radius (Bruker, nominal spring constant 100 pN/nm and 45 kHz resonance frequency in air) were used to investigate cell properties in 2D-gels. MLCT-SPH-5UM (Bruker, nominal spring constant 150 pN/nm and 17 kHz resonance frequency in air) with 5.5 µm tip radii were employed to assess gel properties in 3D-environment (fibroblasts inside the gels). Samples were fixed to an aluminum holder with vacuum grease and mounted on the AFM stage with two magnets. The entire set-up was enclosed in a home built polymethylmethacrylate (PMMA) box in order to inject and maintain 5% CO_2_ during experiments.

Apparent Young’s modulus values were extracted from regular force curves and sweep frequency method was used to obtain cell rheological properties. For measuring cells seeded on 2D-collagen matrices, force map scan size was 5 μm and composed of 16 or 256 force curves (4 x 4 or 16 x 16 lines per frame). Typically, force curves were recorded at a scan rate of 2 Hz; corresponding to a maximum velocity of 20 µm/s. Indentation depths were always greater than 500 nm in order to average the stiffness over a large contact area. For 3D-gels measurements, force curves were acquired over a large area 20 x 20 µm and force maps were composed of 16 or 36 force curves (4 x 4 or 6 x 6 lines per frame). Force curves were recorded at a scan rate of 1 Hz, corresponding to a maximum probing velocity of 7.94 µm/s.

### AFM data analysis

The data analysis software IGOR (Wavemetrics, Lake Oswego, OR, USA) was used to evaluate the mechanical properties of cells in terms of apparent Young’s modulus (*E*). The Hertzian model for spherical tips was used to calculate the apparent Young’s modulus for each force curve within a force map. The median and 25/75 percentiles and logarithmic histogram of apparent Young’s modulus were considered as representative modulus of each force map. Sweep frequency data were fitted with the power law structural damping model. *E\** data are separated into real (in phase) and imaginary (out of phase) parts. The real part represents the storage modulus and it is a measure of the elastic energy stored and recovered per cycle of oscillation. The imaginary part depicts the loss modulus and it accounts for the energy dissipated per cycle of sinusoidal deformation. We also calculate the loss tangent, which is an index of the solid-like (<1) or the liquid-like (>1) behavior of the cell. This model assumes a storage modulus that increases with frequency following a power law with exponent α, and a loss modulus that includes a term that is a fraction η of the storage modulus and a Newtonian viscous term.

### Live/Dead staining

After 2 days, 1 week and 2 weeks of culture, cells were stained to assess cell viability within 3D collagen gels. Live-dead kits (ThermoFisher, R37601) were used by following manufacturer instructions. Calcein AM solution was transferred to BOBO-3 Iodid solution and mixed. All content was added to the 3D-gels with the same amount of cell culture medium. Samples were incubated for 15 minutes at room temperature and finally imaged. Calcein AM provides green color to alive cells and BOBO-3 Iodid red to dead cells. Nikon Eclipse Ti Inverted epifluorescence Microscope (Nikon Instruments Inc., Melville, New York) with a 20x objective lens was used to image the samples.

### Cell staining for morphological analysis

Cells seeded on 2D-collagen gels and petri dishes were fixed with 4% paraformaldehyde for 30 minutes and permeabilized with 0.2% Triton X100 for 5 minutes. Samples were washed with PBS after each step and then incubated with 10% FBS for 30 minutes at room temperature. ActinRed (Invitrogen, ready probes) was used to label actin fibers and NucBlue (Invitrogen, ready probes) for nuclear staining, two drops of each solution was added to 1 mL PBS and then incubated for 30 minutes at room temperature. After three times washing with PBS, cells were finally stored in PBS at 4°C prior to image acquisition. Nikon Eclipse Ti Inverted epifluorescence Microscope (Nikon Instruments Inc., Melville, New York) with a 40x objective lens was used to observe cells and collect fluorescent images.

Eccentricity of the cells was computed by adjusting an ellipse to the polygon described generated by ImageJ software, as described in^51^. Cell number inside each 3D collagen gels was obtained counting the number of nucleus in each z-stack layer using Image J software.

### Gel immunostaining

3D-gels were stained for fibronectin to assess the secretion of ECM proteins by the different fibroblasts. Samples were fixed with 4% PFA for 30 minutes at room temperature. Samples were then blocked with PBS, 0.4 % Triton and 10 % donkey serum solution for one hour at room temperature. Primary antibody against fibronectin (mouse anti-fibronectin, 1:50, ThermoFisher, CSI 005-17-02) was incubated in a solution of PBS, 0.4 % Triton and 0.1 % donkey serum for 24h at room temperature. The samples were rinsed three times with the same solution. Hoechst 33342 (1:1000, A-21202 Invitrogen) was used to visualize cell nucleus and secondary antibody (donkey anti-mouse AlexaFlour 488, 1:1000, H3570 Invitrogen) was incubated in PBS and 0.4 % Triton for 2 h at 37°C. Three times rinsed with PBS were applied to eliminate unbound secondary antibodies. Olympus FluoVIEW 3000 RS confocal microscope with UPLXAPO 10x and 20x objective lens was used to acquire images.

### Statistical analysis

Differences between the apparent Young’s modulus and rheological parameters of different cells or gels were determined with Wilcoxon signed-rank test and Cohen’s d test using IGOR. P-values obtained from Wilcoxon signed-rank test, * indicating p < 0.01 and Cohen’s d test with # indicating 0.2 < d < 0.5 and ## indicating d > 0.5.

## Data availability statement

The datasets generated during and/or analyzed during the current study are available from the corresponding author on reasonable requests.

## Acknowledgements

We would like to thank Cesare Covino for acquiring gel’s immunofluorescence images. We would like to thank Prof. Ursula Mirastschijski for providing tissue samples and Prof. Gazanfer Belge for stabilizing cell lines from primary cells. We would like to thank Nina Messerschmidt for the help in sample preparation. We would also like to thank the School of Medicine at the University of Barcelona for providing rat-tails to obtain collagen. We would also like to thank Prof. Dorothea Brüggemann for the use of the epifluorescence microscope and Dr. Arundhati Joshi and Dr. Deepanjalee Dutta for the helpful discussions and suggestions.

## Funding

This project has received funding from the European Union’s Horizon 2020 research and Innovation programme under the H2020-MSCA-ITN-2018 Grant Agreement No. 812772.

The authors acknowledge Euro-BioImaging (www.eurobioimaging.eu) for providing access to imaging technologies and services via the Italian Node (ALEMBIC, Milan, Italy).

## Author contributions

S.P-D. designed the experimental scheme and performed AFM measurements, data analysis, collagen gels preparation, cell’s fluorescent images in 2D and 3D and manuscript preparation. H.S.F. helped obtaining collagen from rat-tails, preparing collagen gels and manuscript preparation. L.M.V. helped establishing gel’s immunofluorescence protocol, analysis of gel’s immunofluorescence data and manuscript preparation. M.A. helped establishing gel’s immunofluorescence protocol, analysis of gel’s immunofluorescence data and manuscript preparation. J.O. helped obtaining collagen from rat-tail, collagen gel’s protocol and manuscript preparation and M.R. helped designing the experimental scheme, data analysis and manuscript preparation.

## Additional information

**Supplementary information** accompanies this paper at:

**Competing interest:** The authors declare no competing interests.

## Notes

### Competing Interest Statement

The authors have declared no competing interest.

