## Supporting information for "Characterizing the viscoelastic properties of different fibroblasts in 2D and 3D collagen gels"

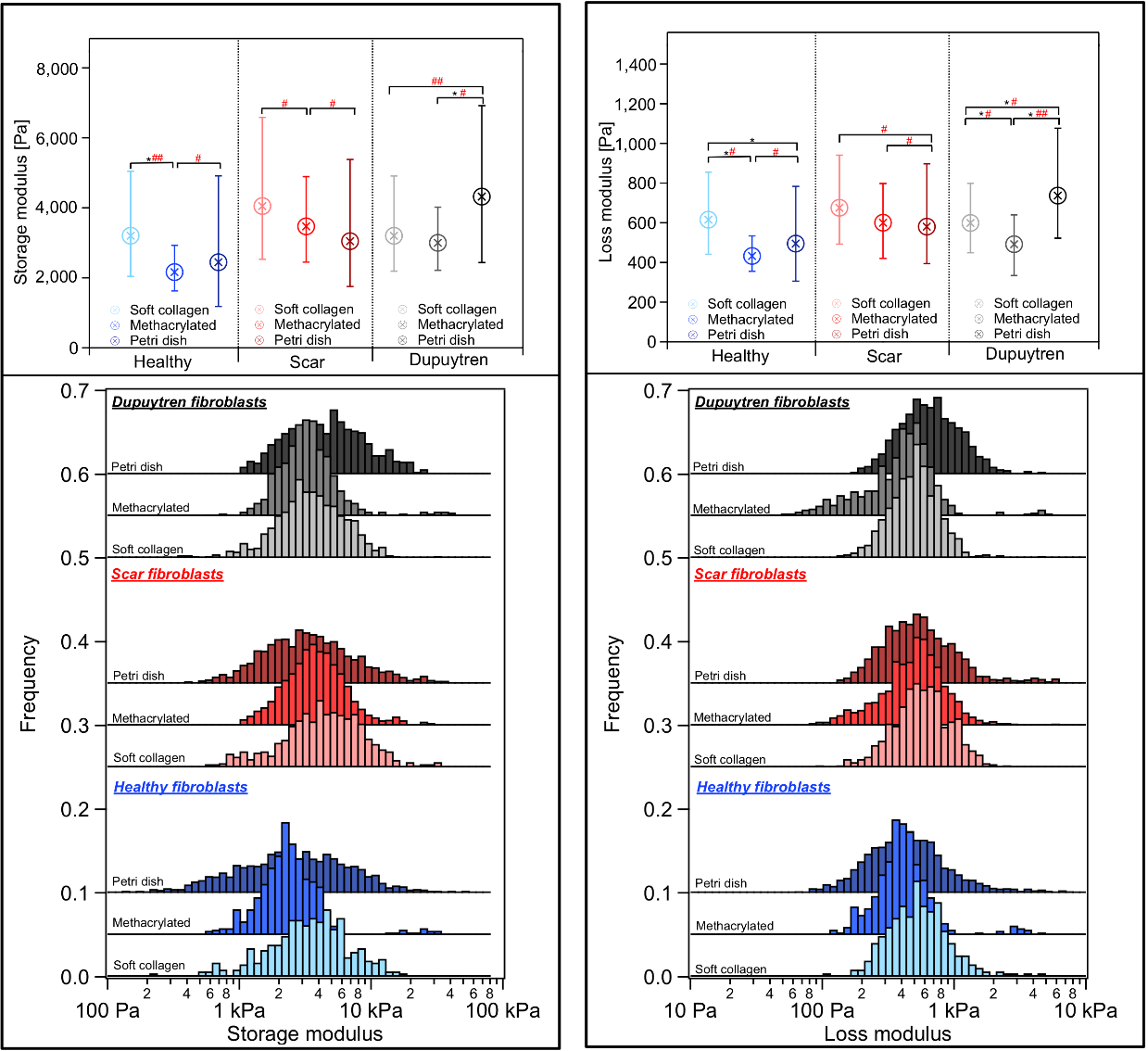

Figure S1 Box plot (median with 25/75 percentiles) and histogram distribution of storage and loss modulus at 1 Hz (*n* = 50). Data are sorted by cell type and within each group each color presents one type of substrate (from lightest to darkest: soft collagen to petri dish).

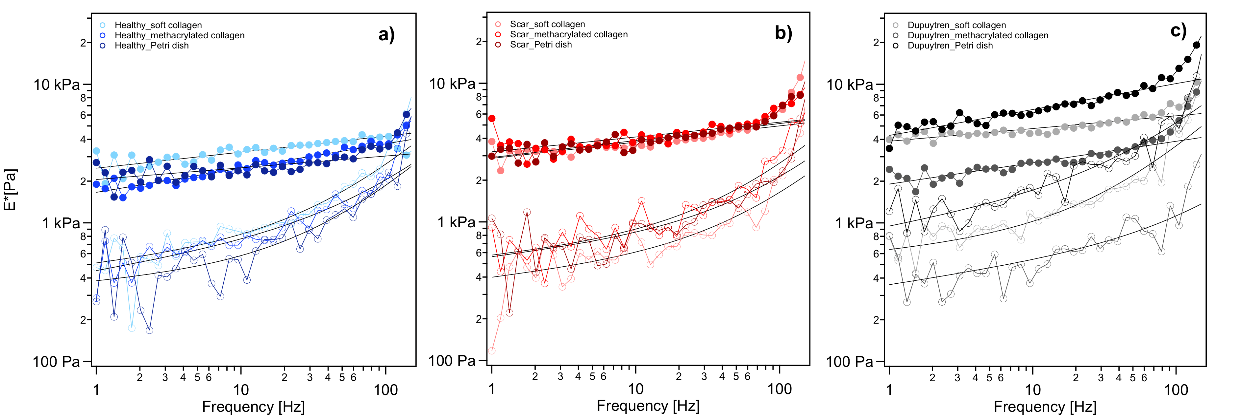

Figure S2 Frequency dependence of storage and loss modulus after hydrodynamic viscous drag correction. Close symbols represent the storage modulus and the open symbols the loss modulus. a) Healthy fibroblasts, b) scar fibroblasts and c) Dupuytren fibroblasts. From lightest to darkest color in each graph represents fibroblasts seeded on soft collagen to Petri dish (*n* = 50).

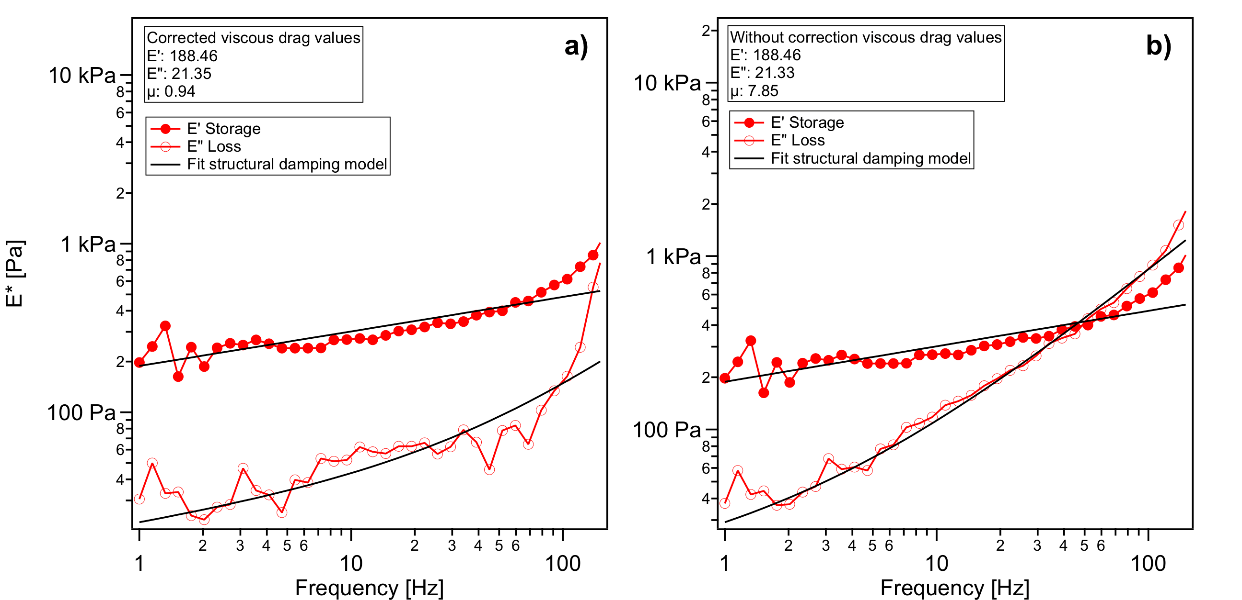

Figure S3 Representation of E* values over frequency, corrected for the viscous drag (a) and without correction (b).

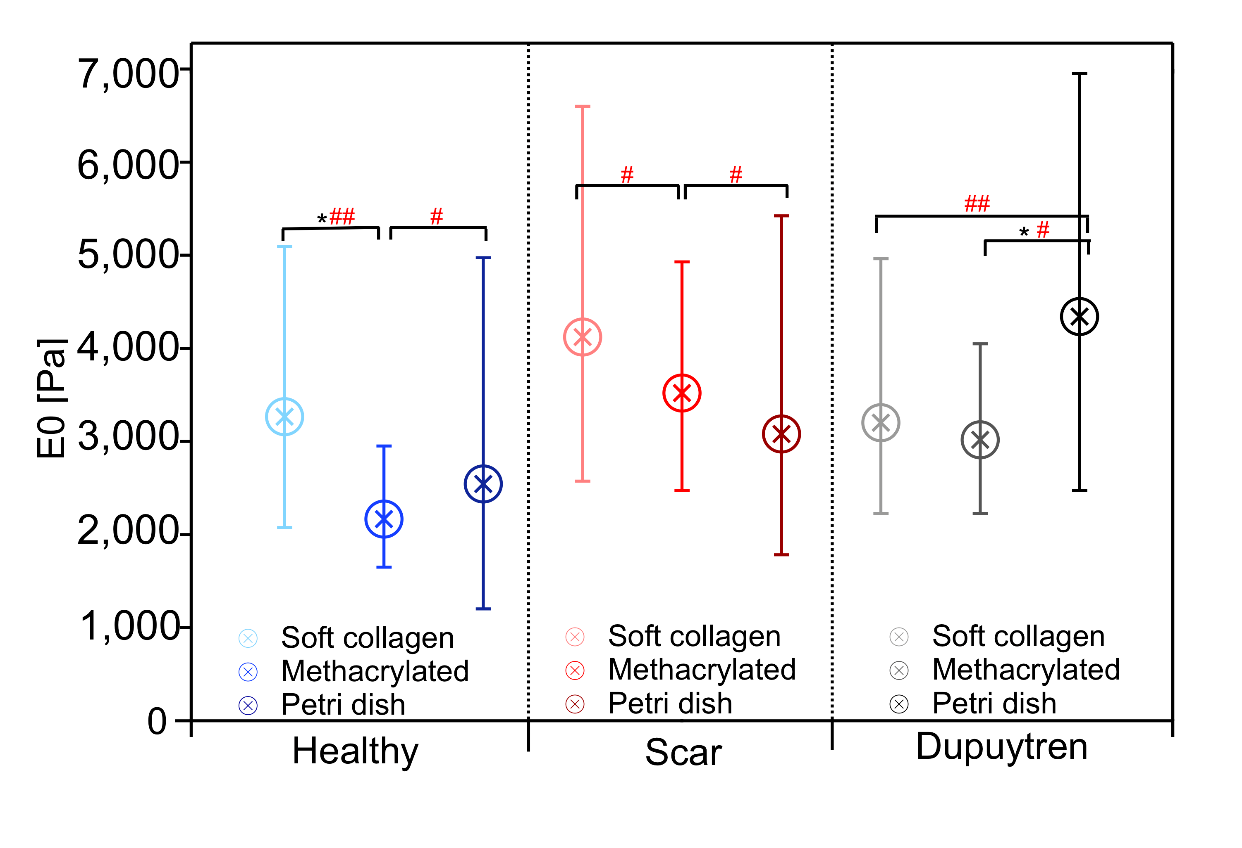
Figure S4 Box plot (median with 25/75 percentiles) of E_0_ (*n* = 50). Data are sorted by cell type and within each group each color presents one type of substrate (from lightest to darkest: soft collagen to petri dish).

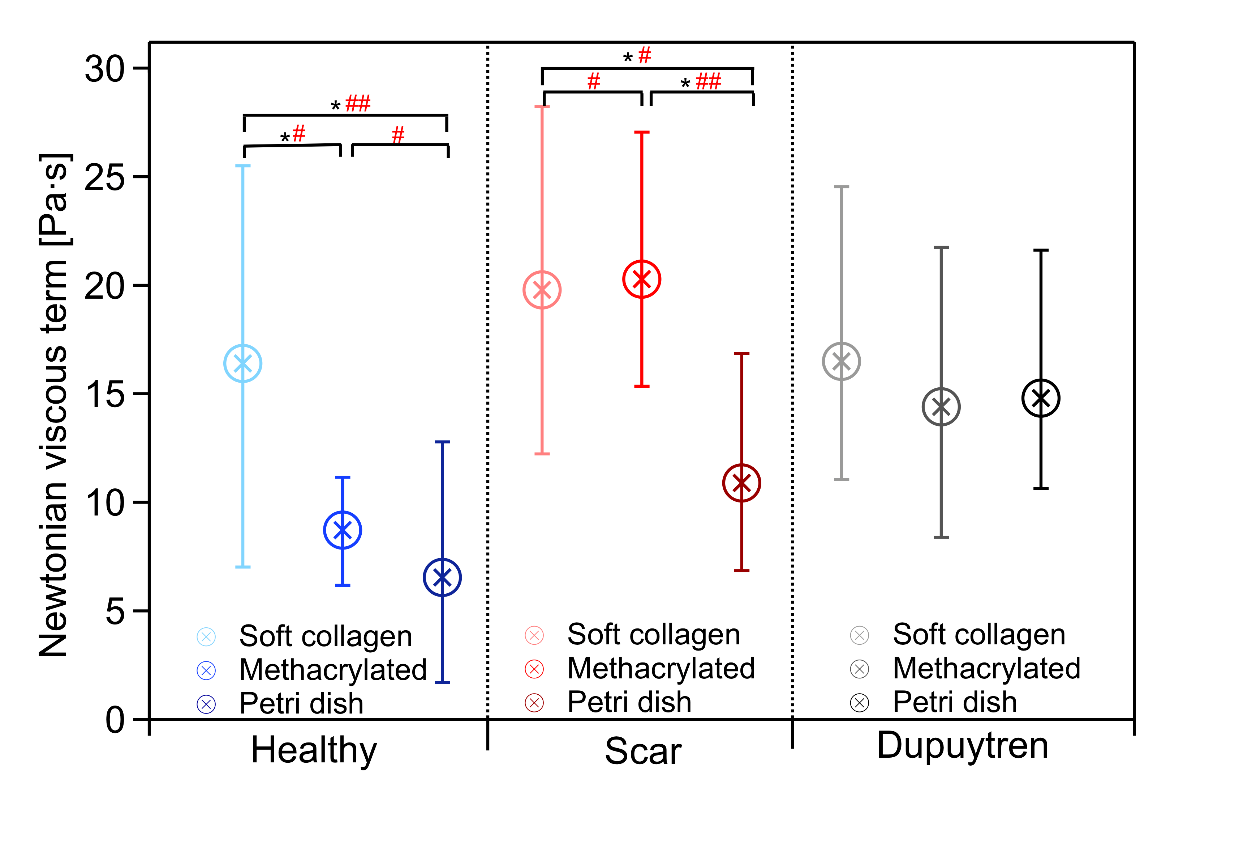

Figure S5 Box plot (median with 25/75 percentiles) of *μ* (*n* = 50). Data are sorted by cell type and within each group each color presents one type of substrate (from lightest to darkest: soft collagen to petri dish).

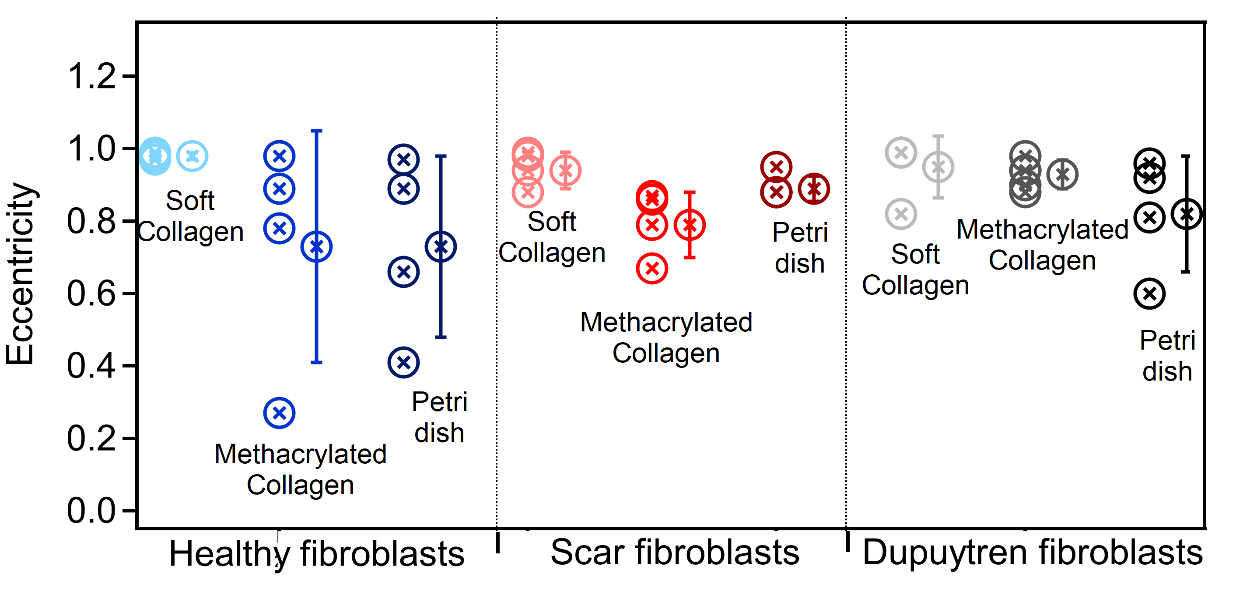

**Figure S6** Quantification of cell eccentricity to characterize their morphology. Mean ± standard deviation; symbols with the same color next to the mean value correspond to the individual measurements. Data are sorted by cell type and within each group each color presents one type of substrate (from lightest to darkest: soft collagen to petri dish).

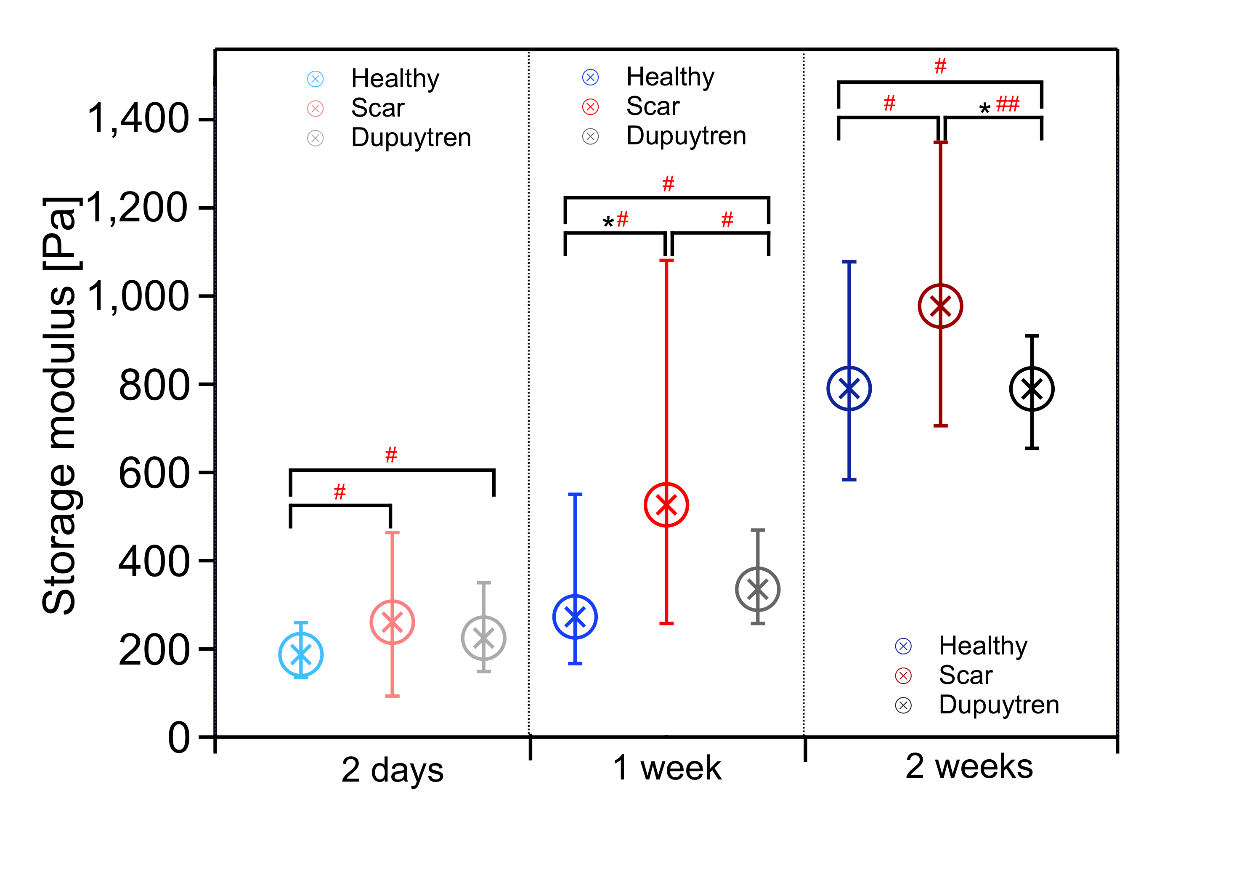

Figure S7 Box plot (median with 25/75 percentiles) of E’, storage modulus at 1 Hz (*n* = 30). Within each group (incubation time) each color presents one type of fibroblasts (blue healthy, red scar and black Dupuytren fibroblasts).

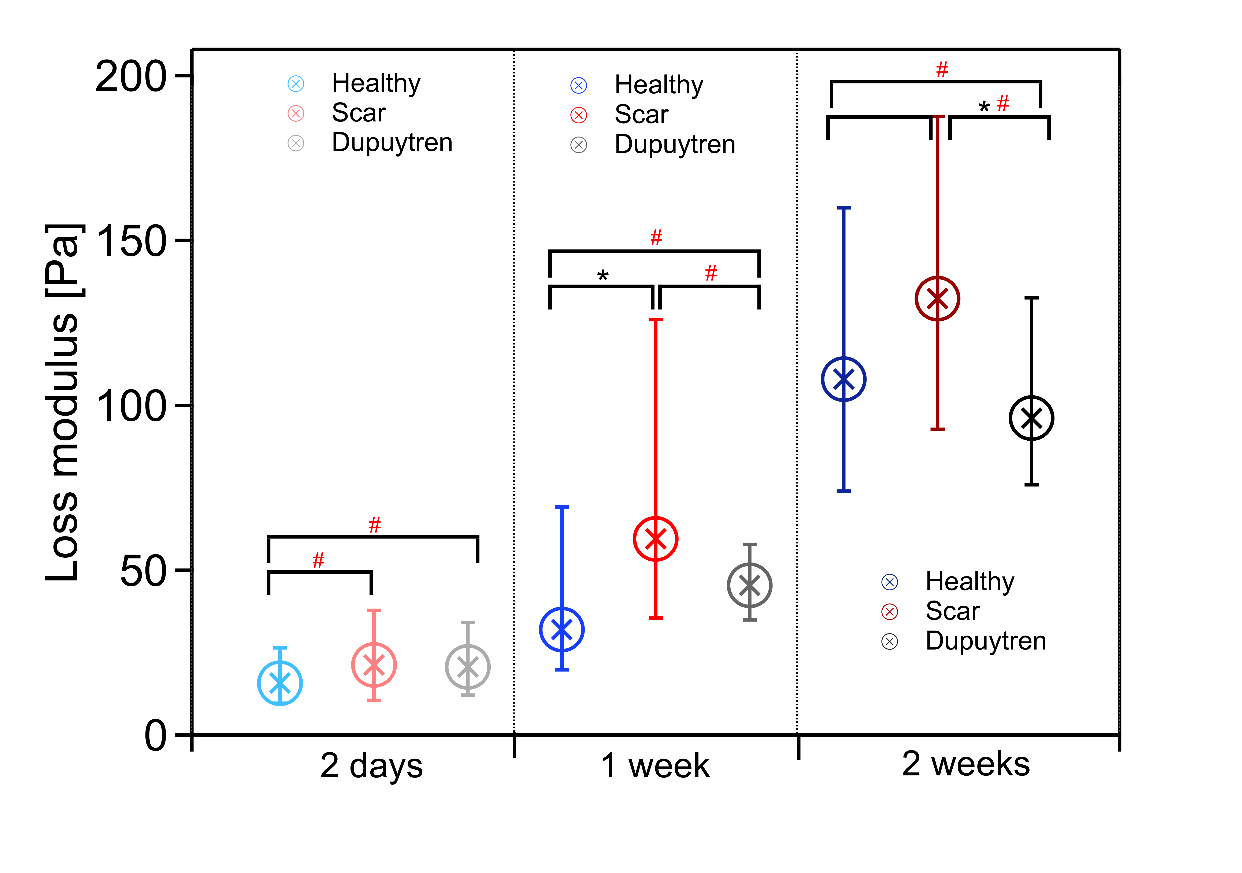

Figure S8 Box plot (median with 25/75 percentiles) of E”, loss modulus at 1 Hz (*n* = 30). Within each group (incubation time) each color presents one type of fibroblasts (blue healthy, red scar and black Dupuytren fibroblasts).

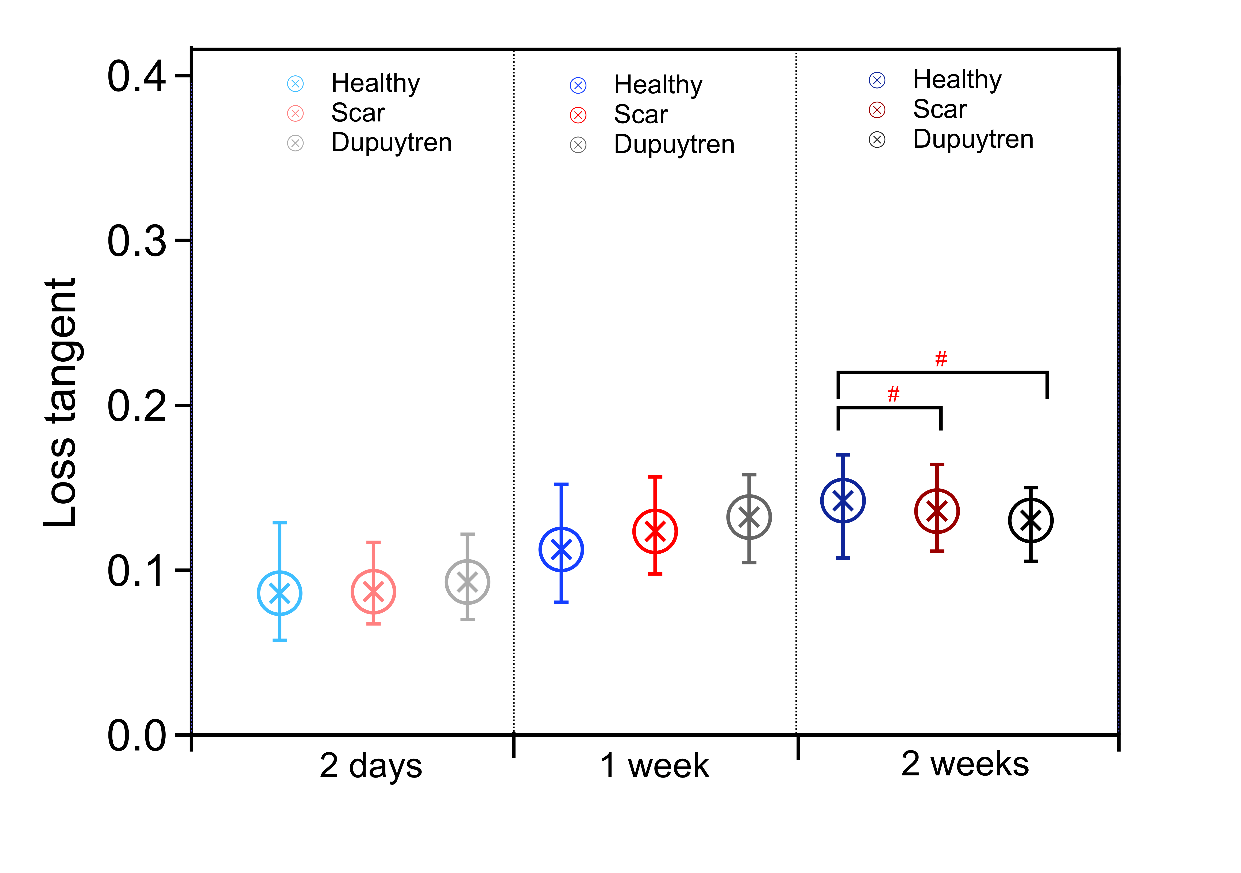
Figure S9 Box plot (median with 25/75 percentiles) of η, loss tangent at 1 Hz (*n* = 30). Within each group (incubation time) each color presents one type of fibroblasts (blue healthy, red scar and black Dupuytren fibroblasts).

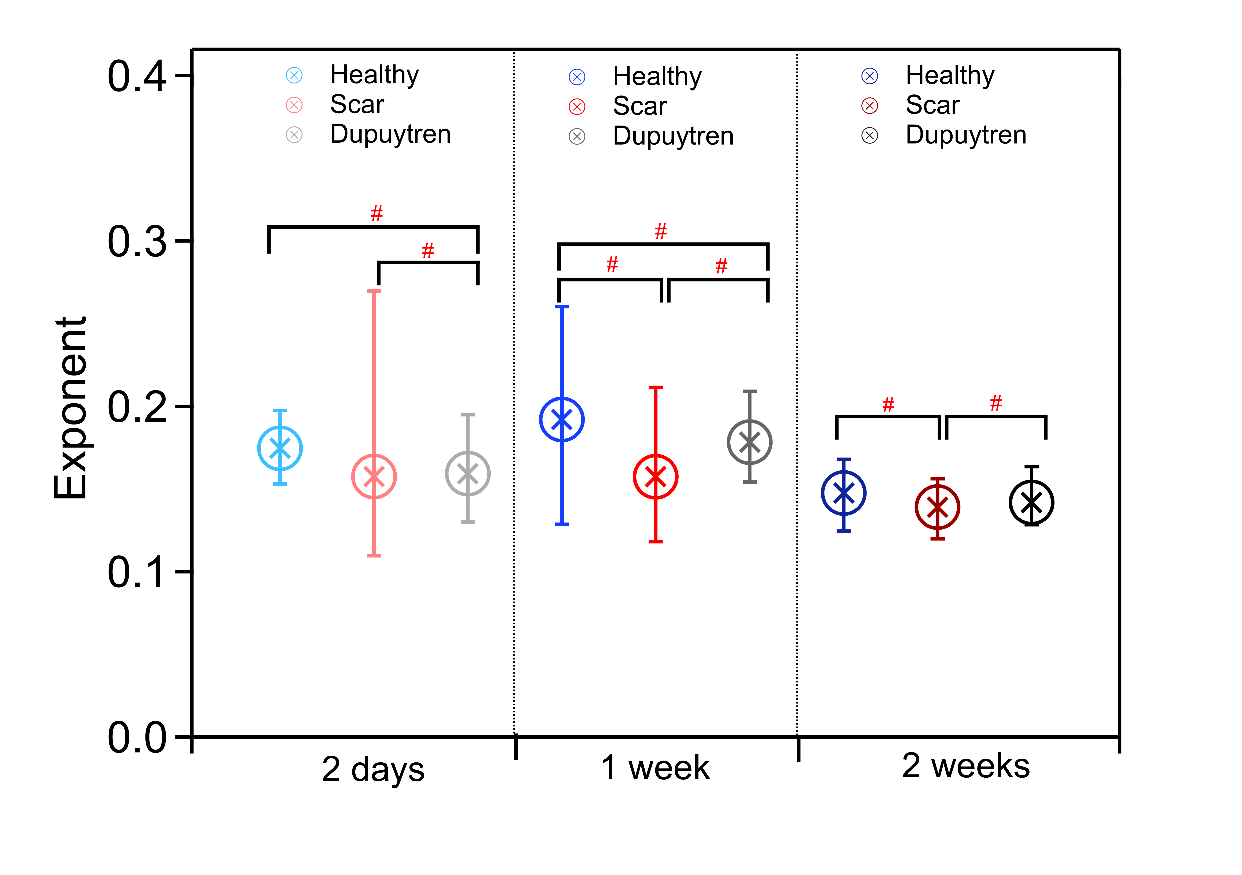

Figure S10 Box plot (median with 25/75 percentiles) of α, exponent (*n* = 30). Within each group (incubation time) each color presents one type of fibroblasts (blue healthy, red scar and black Dupuytren fibroblasts).

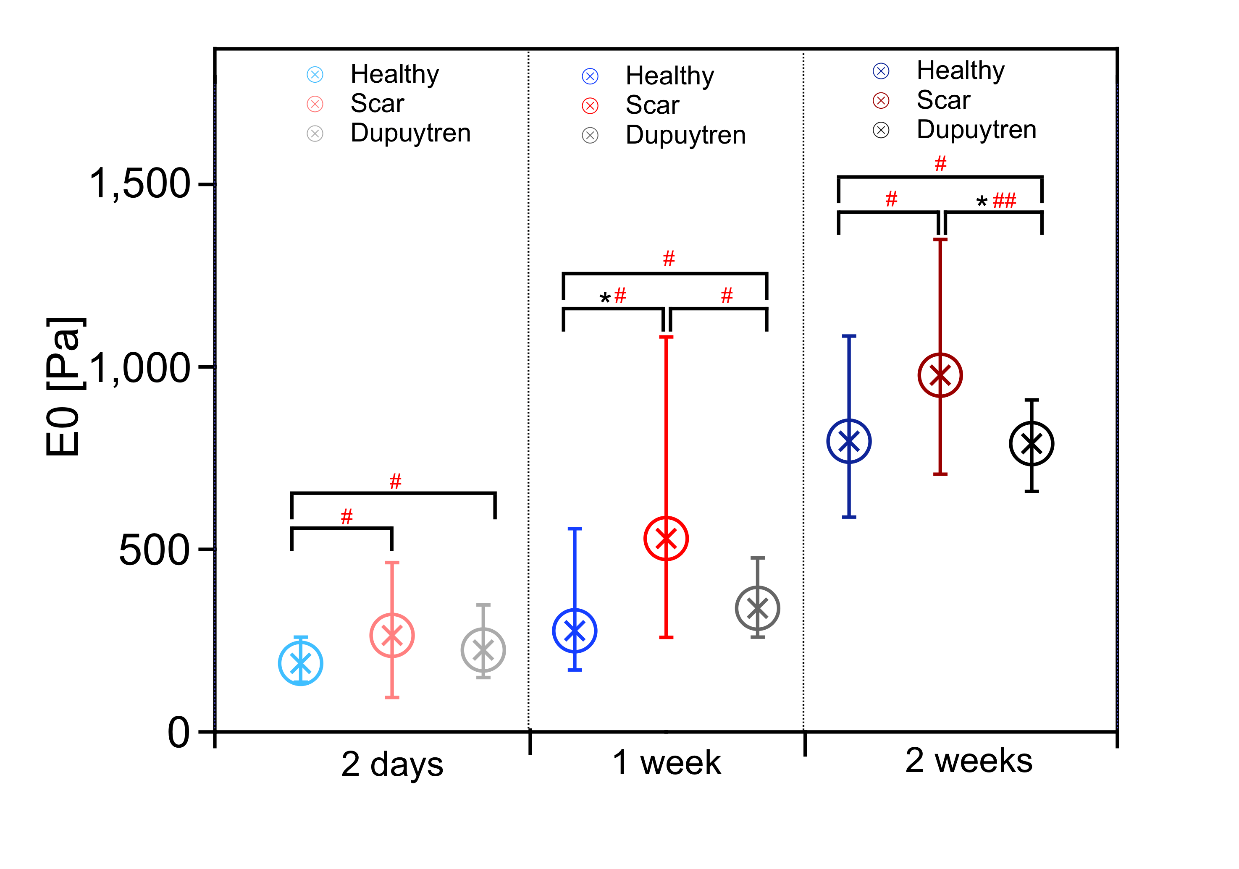

Figure S11 Box plot (median with 25/75 percentiles) of E_0_ (*n* = 30). Within each group (incubation time) each color presents one type of fibroblasts (blue healthy, red scar and black Dupuytren fibroblasts).

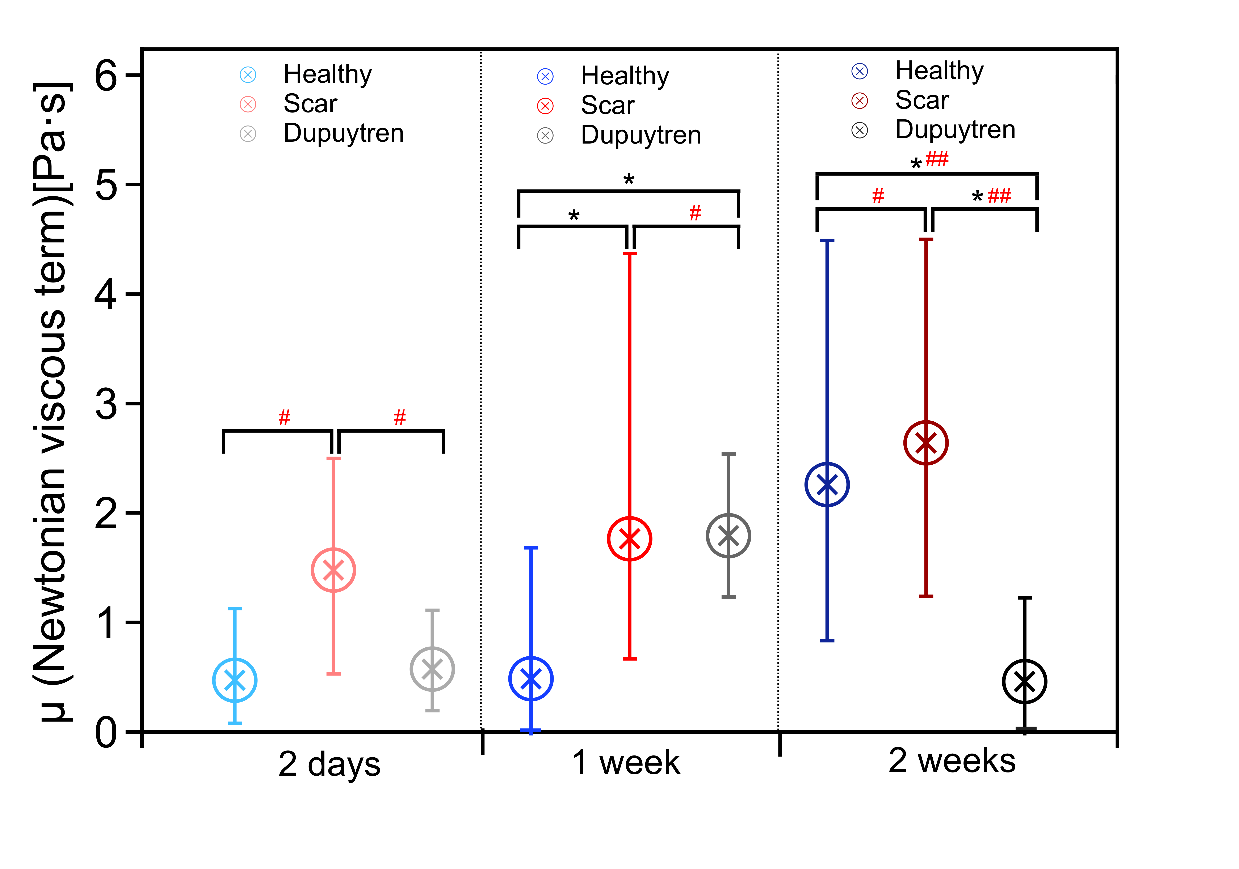

Figure S12 Box plot (median with 25/75 percentiles) of *μ* (*n* = 30). Within each group (incubation time) each color presents one type of fibroblasts (blue healthy, red scar and black Dupuytren fibroblasts).

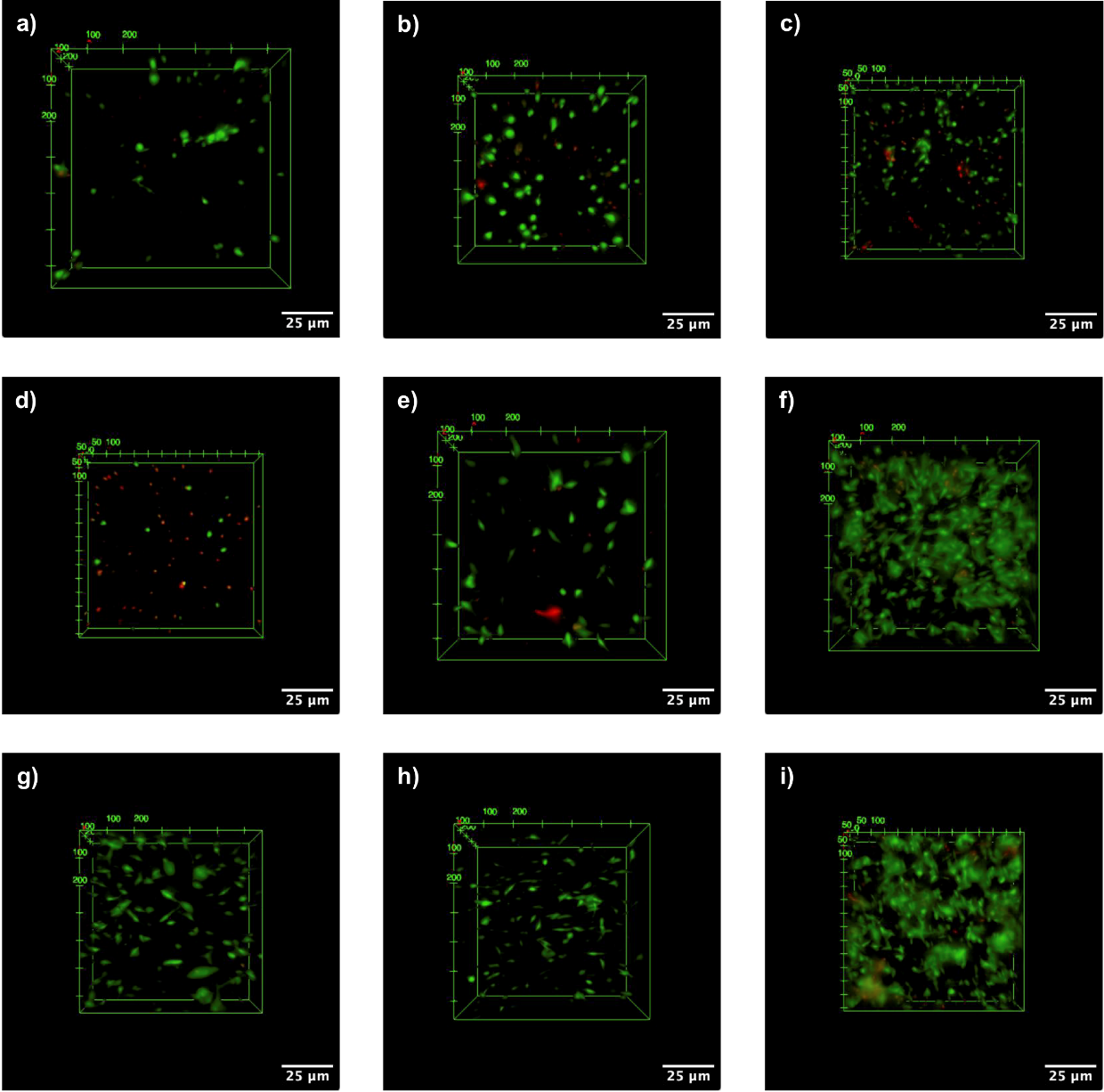

Figure S13 Live/dead staining of fibroblasts in all gel conditions. Alive cells are labelled in green and dead cells in red. 2 days incubation: a) healthy, b) scar and c) Dupuytren fibroblasts; 1 week incubation: d) healthy, e) scar and f) Dupuytren fibroblasts and 2 weeks incubation: g) healthy, h) scar and i) Dupuytren fibroblasts. Scale bar: 25 µm.

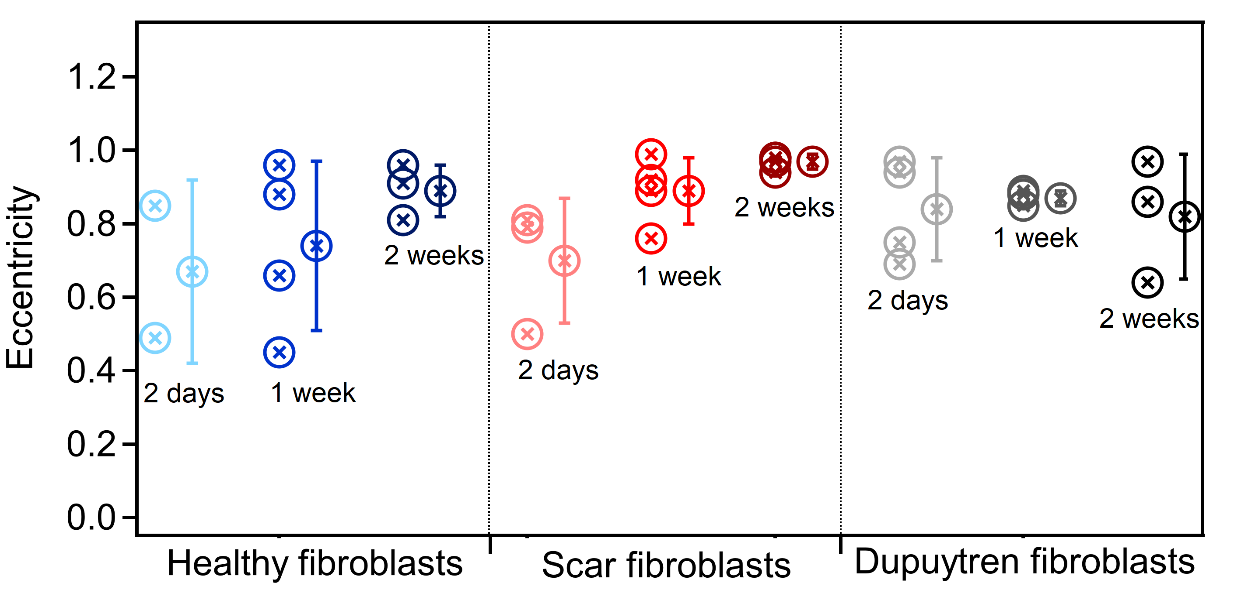

**Figure S14** Quantification of cell eccentricity to characterize their morphology. Mean ± standard deviation; symbols with the same color next to the mean value correspond to the individual measurements.
